# High levels of endemism and local differentiation in the fungal and algal symbionts of saxicolous lecideoid lichens along a latitudinal gradient in southern South America

**DOI:** 10.1101/699942

**Authors:** Ulrike Ruprecht, Fernando Fernández-Mendoza, Roman Türk, Alan Fryday

## Abstract

Saxicolous, lecideoid lichenized-fungi have a cosmopolitan distribution but, being mostly cold adapted, are especially abundant in polar and high-mountain regions. To date, little is known of their origin or the extent of their trans-equatorial dispersal. Several mycobiont genera and species are thought to be restricted to either the northern or southern hemisphere, whereas others are thought to be widely distributed and occur in both hemispheres. However, these assumptions often rely on morphological analyses and lack supporting molecular genetic data. Also unknown is the extent of regional differentiation in the southern Polar Regions.

An extensive set of lecideoid lichens (185) was collected along a latitudinal gradient at the southern end of South America, always staying in areas of subantarctic climate by increasing the elevation of the collecting sites with decreasing latitude. The investigated specimens were brought into a global context by including Antarctic and cosmopolitan sequences from other studies. For each symbiont three markers were used to identify intraspecific variation (mycobiont: ITS, mtSSU, RPB1; photobiont: ITS, psbJ-L, COX2). For the mycobiont the saxicolous genera *Lecidea, Porpidia, Poeltidea* and *Lecidella* and their photobionts *Asterochloris* and *Trebouxia* were phylogenetically revised. The results show for several globally distributed species groups geographically highly differentiated subclades, classified as operational taxonomical units (OTUs), which were assigned to the different regions of southern South America (sSA). Further, for sSA, several small endemic and well supported clades were detected at the species level for both symbionts.

## Introduction

Saxicolous ‘lecideoid’ lichens (Hertel 1984) comprise genera and species described under the generic name *Lecidea* sensu Zahlbruckner (1925) and comprise crustose species with apothecia lacking a thalline margin and with non-sepate ascospores. As such they are a heterogenous, non-monophyletic group and, although some do belong to the genus *Lecidea* s. str. (Lecideaceae), most belong to other genera and families, such as *Carbonea, Lecidella* (Lecanoraceae), *Porpidia*, *Poeltidea* and *Cyclohymenia* (Lecideaceae). In addition to their morphological similarities, lecideoid lichenized fungi are strongly associated with green microalgal photobionts of the cosmopolitan class Trebouxiophyceae (Buschbom and Mueller 2004; Fryday and Hertel 2014; Hertel 1984; 2007; Ruprecht et al. 2012a; Ruprecht et al. 2016; Schmull et al. 2011; Zhao et al. 2015).

The inconspicuous morphology of this group complicates their systematic treatment. Large taxonomic groups are often distinguishable by just a few microscopic traits, such as spore size and septation or ascus-type, but species level identification can be difficult, often relying on cryptic characters like excipulum pigmentation and structure or the secondary metabolites produced. The fundamental taxonomic work on lecideoid lichens (Castello 2003; Fryday 2005; Fryday and Hertel 2014; Gowan 1989; Hafellner 1984; Hertel 1984; 1995; Inoue 1995; Knoph and Leuckert 1994; Knoph et al. 2004) mostly used morphological and chemical characters, but lacked molecular genetic data. Extensive collections, especially from the southern hemisphere are very often older than 50 years, which precludes the use of molecular methods, because of the common problem of DNA degradation in mycobiont specimen older than 20 years. During the last decade, molecular re-evaluations have helped to redefine the species concepts behind these diverse groups but were mostly focused on the northern hemisphere (Buschbom and Mueller 2004; McCune et al. 2017; Orange 2014; Schmull et al. 2011; Zhao et al. 2015; Zhao et al. 2016) and Antarctica (Ruprecht et al. 2010; 2012b), although the links to a better understanding of distribution- and speciation patterns in the group are still largely missing. Meanwhile, intermediate latitudes in the southern hemisphere remain understudied and recently published results (Ruprecht et al. 2016) have emphasized the extent of the knowledge gap in southern South American lecideoid lichens, not only from the mycobiont perspective but also from that of the associated green microalgae.

The use of DNA sequence data and phylogenetic methods has largely contributed to the idea that cosmopolitan genera often show locally differentiated subgroups or cryptic species, which can be influenced by ecological factors and may be restricted to isolated areas (Branco et al. 2015; Kraichak et al. 2015; Leavitt et al. 2011; Lumbsch and Leavitt 2011; Walser et al. 2005). Lichens, as well as non-lichenized fungi, with an arctic-alpine distribution in the northern hemisphere are, however, a notable exception to this pattern, often comprising relatively homogenous genetic entities, mostly at the species level, with widespread distributions. A number of studies, such as those on *Porpidia flavicunda* (Buschbom 2007), *Flavocetraria cucullata* and *F. nivalis* (Geml et al. 2010) as well as for several different types of fungi (Geml 2011), indicate continuing intercontinental gene-flow in species that are present in both the northern and southern hemispheres. However, trans-equatorial dispersal is less frequently shown for other, similar lineages, such as the lichenized-fungal genus *Lichenomphalia* (Geml et al. 2012) or the species *Cetraria aculeata* (Fernandez-Mendoza and Printzen 2013). In contrast, although the distribution of green algal photobionts of the genus *Trebouxia* extends across broad intercontinental regions, especially in the northern hemisphere (Leavitt et al. 2015), a pattern of trans-equatorial dispersal with low diversification is common (Muggia et al. 2010; Ruprecht et al. 2012a). However, for the mycobionts, strong diversification and endemism in the southern hemisphere is expected, resulting in the development of several distinct species and genera; for example, *Lecidea aurantia, L. cambellensis* (Fryday and Hertel 2014) or *L. cancriformis* (Castello 2003), *Poeltidea* (Hertel 1984), *Gondwania, Shackletonia* (Arup et al. 2013) and *Protousnea* (Calvelo et al. 2005). Interestingly, this does not seem to be the case for the cosmopolitan green algal genus *Trebouxia*, although one exception was found in the most extreme areas in continental Antarctica (Ruprecht et al. 2012a).

The most probable scenarios for disjunct distributions is that they can be caused (1) by vicariant history or long distance dispersal, such as for the species *Staurolemma omphalarioides*, which is known from the Mediterranean region and Norway (Bendiksby et al. 2014), (2) by transition from the Arctic to Patagonia in the Pleistocene and resulting in cryptic specialisation as shown in the bipolar lichen *Cetraria aculeata* (Fernandez-Mendoza and Printzen 2013) and (3) by glacial refugia during the last ice ages at the southern end of South America (Paula and Leonardo 2006). A good example of this is the highly differentiated and endemic lichen species *Porpidia navarina,* which is known only from one of the southernmost islands (Isla Navarino) that was ice free during the Last Glacial Maximum (Douglass et al. 2005; Ruprecht et al. 2016). (4) Finally, adaptation and subsequent specialisation to the harsh climate conditions in Antarctic cold deserts (Ruprecht et al. 2012a; Ruprecht et al. 2010; Schroeter et al. 2011) can also lead to the high local differentiation in global species and endemism in the southern Polar regions.

Lichens are ideal model-systems to test these hypotheses, because several genera and species are globally distributed and form locally differentiated subgroups (Fernandez-Mendoza et al. 2011). Additionally, at least double the information is available compared to other organisms because lichens consist of a symbiotic relationship of two or more independently distributed partners. This main symbiotic relationship is formed by a fungus (mycobiont) and green algae and/or cyanobacteria (photobiont). Additionally, a diverse community of associated bacteria (Aschenbrenner et al. 2016; Grube et al. 2015), algae (Moya et al. 2017; Peksa and Skaloud 2011; Ruprecht et al. 2014), endolichenic or lichenicholous fungi and basidiomycete yeasts (Arnold et al. 2009; Lawrey and Diederich 2003; Spribille et al. 2016) are part of the lichen thallus.

This study focuses on the geographically isolated, tapering southern end of the South American continent (southern Patagonia, including the islands around Tierra del Fuego and Cape Horn). Due to climatic conditions equivalent to Maritime Antarctica, the southern subpolar region (or subantarctic subregion), which is characterized by an absence of arboreal vegetation (Brummitt 2001; Morrone 2000) is included as part of the Antarctic floral kingdom (Takhtajan and Cronquist 1986). The subantarctic subregion extends northwards through the continent at increasing elevations along the mountain ranges of the southern Andes (Morrone 2000). To the south, Maritime Antarctica is the closest landmass, separated by about 900 km of ocean and the Antarctic Circumpolar Current (Allison et al. 2010; McCave et al. 2014). These areas are colonized by specialized cold-adapted organisms, which often act as pioneer vegetation (Bilovitz et al. 2015; Caccianiga and Andreis 2004; Hertel 1984). Among other organisms, the globally distributed saxicolous lecideoid lichens are one of the most frequent vegetation types, forming diverse communities on rocks and boulders, mainly in treeless areas above a temperate rainforest (Fig. 1; Goffinet et al. 2012; Hertel 2007; Ruprecht et al. 2016).

**Figure 1:**
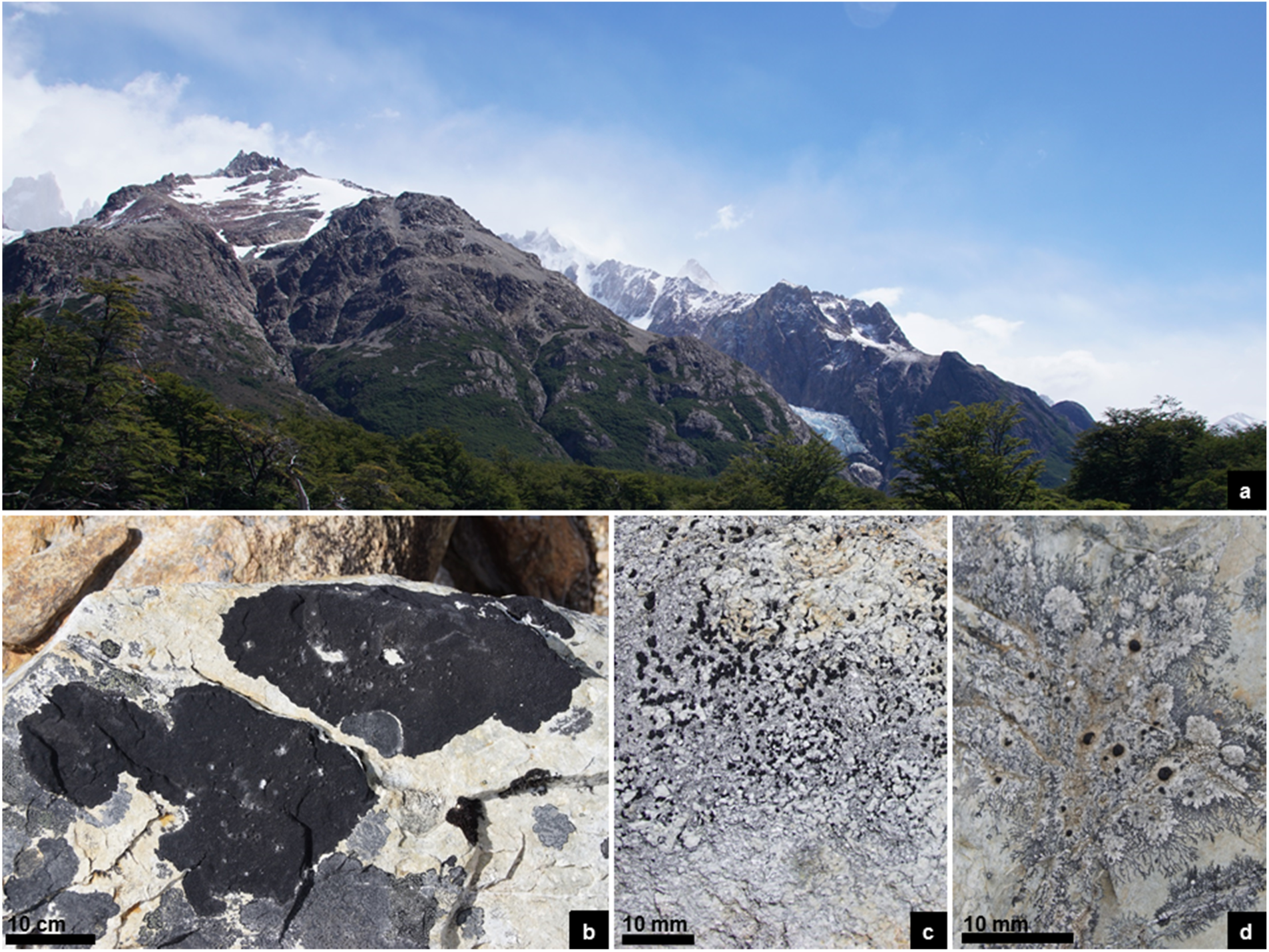
a) Classical subantarctic subregion above tree level: Parque Nacional Los Glaciares, Argentina; (b) saxicolous crustose lichens on siliceous rock: *Lecidea auriculata, L. kalbii, Poeltidea perusta, Rhizocarpon geographicum;* (c) *Lecidea lapicida;* (d) inc. sed. *Poeltidea* sp. 1.

In the current study, the generic phylogenetic classification of the two dominant symbionts (myco- and photobiont), along with their distribution and diversification along climate gradients, were re-evaluated with three marker datasets for each symbiont based on several phylogenetic methods (distance, model based and bGMYC). To accomplish this, both symbionts of saxicolous lecideoid lichen specimens from sSA were placed in a global context using sequence information from the first author’s project framework, from collaborating partners and from official databases (e.g. GenBank). Additionally, newly generated sequences of species that had previously only been described morphologically were included, as well as sequences of uncertain application in the genera *Lecidea* s. str. (Hertel 1984), *Porpidia, Poeltidea*, *Lecidella* (mycobiont) and *Asterochloris* and *Trebouxia* (photobiont).

## Material and Methods

### Collecting sites and material

One hundred and eighty five saxicolous lecideoid lichen samples were collected in southern South America along a latitudinal gradient following the subantarctic climatic subregion by increasing elevation from south (Cerro Bandera, Isla Navarino, CL, S55°, 620 m a.s.l.) to north (Cerro Catedral, Bariloche, AR, S41°, 2100 m a.s.l.) and including some areas at a lower elevation in southern Chile. The specimens were collected from siliceous rock in areas above the treeline that were dominated by subantarctic climatic conditions with an annual mean temperature (BIO_1_) of 0 to 7.8 °C and an annual precipitation (BIO_12_) of 320 to 1640 mm (Karger et al. 2017; Fig. 2, Table S1 & S2).

**Figure 2:**
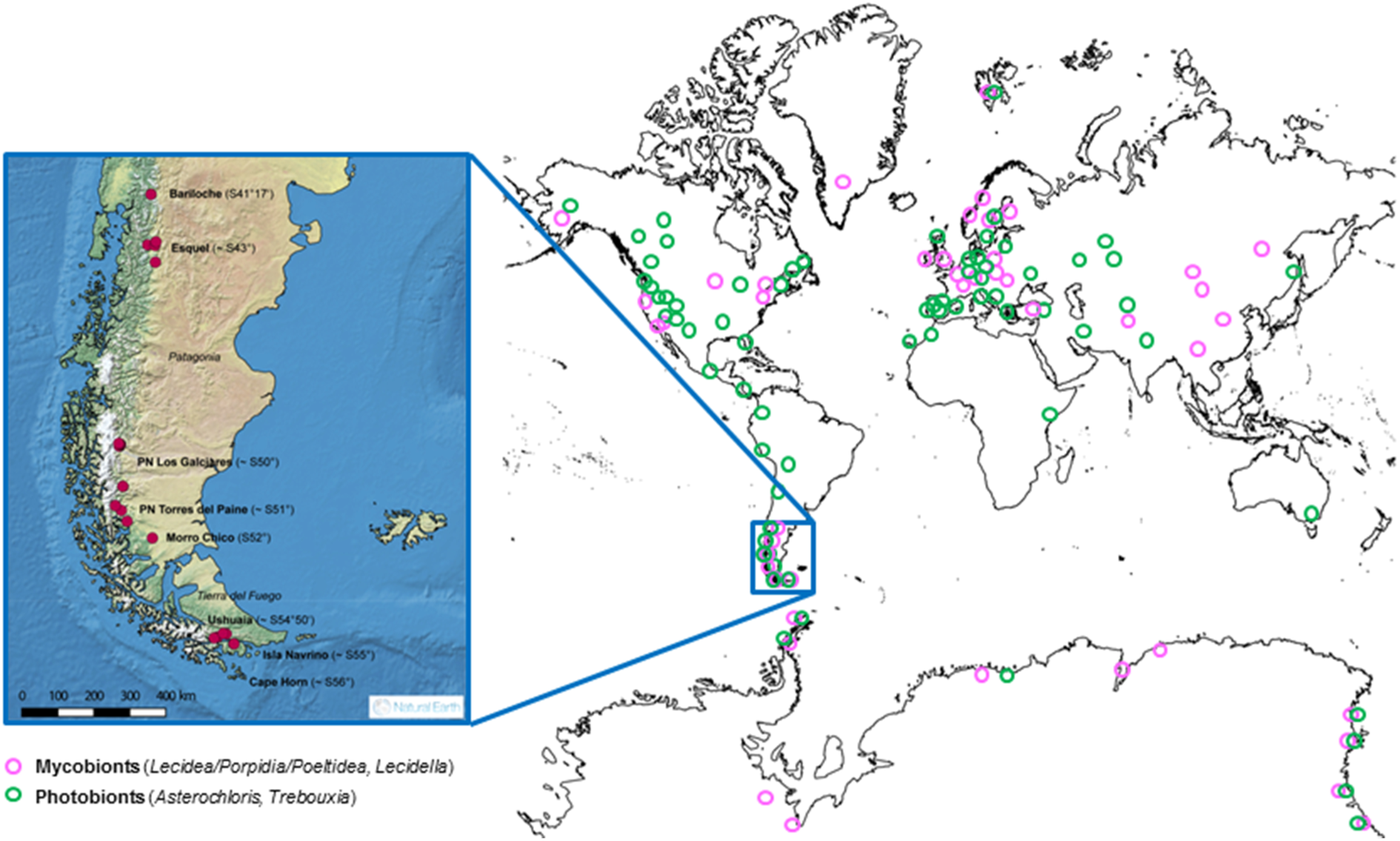
World map showing the locations of the included accessions obtained from Genbank and our own database. Pink circles show the collecting points of the mycobiont and green circles of the photobiont accessions. The enlarged map in color shows the sampling sites from this study in southern South America.

All our specimens collected from southern South America (sSA) are deposited in the Herbarium of the University of Salzburg (SZU).

### DNA amplification and sequencing

Total DNA was extracted from individual thalli by using the DNeasy Plant Mini Kit (Qiagen) according to the manufacturer’s instructions. The PCR mix contained 0.5 units of GoTaq DNA polymerase, 0.2 nM of each of the four dNTPs, 0.3 µM of each primer and about 1 ng genomic DNA.

For each symbiont, three markers were amplified and sequenced with the primers presented in Table S3 with conditions as described in Ruprecht et al. (2014) and Ruprecht et al. (2016). Unpurified PCR-products were sent to Eurofins Genomics/Germany for sequencing.

#### Mycobiont

The internal transcribed spacer region of the nuclear ribosomal DNA (ITS) was amplified for all specimens. Furthermore, the mitochondrial small subunit (mtSSU) and the large subunit of DNA-dependent RNA polymerase 2 (RPB1) were amplified for the *Lecidea/Porpidida/Poeltidea* group.

#### Photobiont

To get a first overview of the different taxa at species and/or genus level of the associated green microalgeae available to the mycobiont, a broad screening along the internal transcribed spacer region of the nuclear ribosomal DNA (ITS) was performed as described in Ruprecht et al. (2016). Additionally, for *Trebouxia* a chloroplast marker, a variable protein-coding gene including an intergenic spacer region (psbJ-L), and a fragment of the mitochondrial cytochrome oxidase subunit 2 gene (COX2) were chosen. Additionally, all three markers were sequenced from four *Trebouxia* cultures (2×S02/Antarctica, A02/Antarctica and A12/Sweden; Table S2d) provided by S. Ott (Düsseldorf), to calibrate the concatenated dataset.

### Phylogenic analyses

Sequences were assembled and edited using Geneious Pro 6.1.8 (www.geneious.com), and aligned with MAFFT v7.017 (Katoh et al. 2002) using preset settings (algorithm: auto select, scoring matrix: 200PAM/k=2 and gap open penalty: 1.34 – 0.123).

Maximum likelihood analyses (ML) were performed using the program IQ-TREE web server (Trifinopoulos et al. 2016) with default settings (Ultrafast bootstrap analyses, 1000 BT alignments, 1000 max. iterations, min correlation coefficient: 0.99, SH-aLRT branch test with 1000 replicates) and presented as a consensus tree. The respective evolutionary models were selected with the implemented model finder (Kalyaanamoorthy et al. 2017) of the program IQ-TREE (Table S4).

The Bayesian phylogenies were inferred using the Markov Chain Monte Carlo (MCMC) procedure as implemented in the program MrBayes 3.2. (Ronquist and Huelsenbeck 2003). The analysis was performed assuming the general time reversible model of nucleotide substitution including estimation of invariant sites and a discrete gamma distribution with six rate categories (GTR + I + Γ; Rodriguez et al. 1990). Two runs with 5 million generations each starting with a random tree and employing 4 simultaneous chains were executed. Every 1,000th tree was saved into a file. Subsequently, the first 25% of trees were deleted as the “burn in” of the chain. A consensus topology with posterior probabilities for each clade was calculated from the remaining 3,751 trees. The phylogenies of the mycobiont of the *Lecidea/Porpidida/Poeltidea* group were rooted with *Farnoldia jurana* subsp. *jurana*, and for *Lecidella* with species of the closely related genera *Lecanora*, *Rhizoplaca* and *Carbonea,* whereas the algal phylogenies were midpoint rooted. All phylogenies were visualized with the program Figtree *v1.4.3* (Rambaut 2014).

### OTU- and cluster delimitation

For each phylogenetically coherent group (*Lecidea/Porpidia/Poeltidea, Lecidella, Asterochloris* and *Trebouxia*) the ITS marker was used to generate OTUs using ABGD (Puillandre et al. 2012). The default settings were used, except ‘X’ (relative gap) at 0.9 and the distance JC69 were chosen. OTUs with a sequence similarity lower than 97.5% were divided to subunits using phylogenetic criteria based on Leavitt et al. (2015) for *Trebouxia* (Table S5).

An alternative cluster delimitation was done by using GMYC models (Reid and Carstens 2012) on the ITS phylogeny *Lecidea/Porpidia/Poeltidea*. Time calibrated phylogenetic reconstructions were carried out in Beast2 (Bouckaert et al. 2014) using the same model settings as for the Bayesian analyses (GTR + I + Γ; Rodriguez et al. 1990) as starting conditions. The suitability of a molecular clock was assessed using the tests implemented in MEGA. The bGMYC method was carried out using the most conservative approach, enforcing a strict clock and a constant size single population prior, thus decreasing the chances of introducing spurious clusters caused by an overfitting the phylogenetic reconstructions.

A maximum clade credibility tree was calculated in treeannotator (Bouckaert et al. 2104) using substitute by: most recent common ancestor (mrca) weights. We used a recursive multi-tree approach implemented in the R package bGMYC (Reid & Carstens 2012) to incorporate phylogenetic uncertainty. The GMYC analysis was iteratively run on a subset of 200 randomly chosen trees using a chain length of 50,000 sampling steps, a burn-in of 40000 and a thinning parameter of 100. The results of all GMYC analyses are summarized in a matrix of pairwise co-assignment probabilities for each haplotype. To obtain a consensus partition making use of k-medioids clustering (Kaufman and Rousseeuw 1990) and optimum average silhouette width to estimate the optimum number of clusters as used in (Ortiz-Alvarez et al. 2015). For the latter we used function *pamk* as implemented in R package *fpc* (Henning 2014) on the co-assignment matrix converted into its dissimilarity correlate.

## Results

### Phylogenetic analyses, OTU and cluster delimitation

Four overall phylogenies for the saxicolous genera *Lecidea/Porpidia/Poeltidea* (Lecideaceae), *Lecidella* (Lecanoraceae), *Asterochloris* and *Trebouxia* (Trebouxiophyceae) using the marker ITS were calculated (Figs. 3 - 6). Additionally, two multi-marker trees for *Lecidea/Porpidia/Poeltidea* (ITS/mtSSU/RPB1; Fig. S1) and *Trebouxia* (ITS/psbJ-L/COX2; Fig. S2) were also calculated.

**Figure 3:**
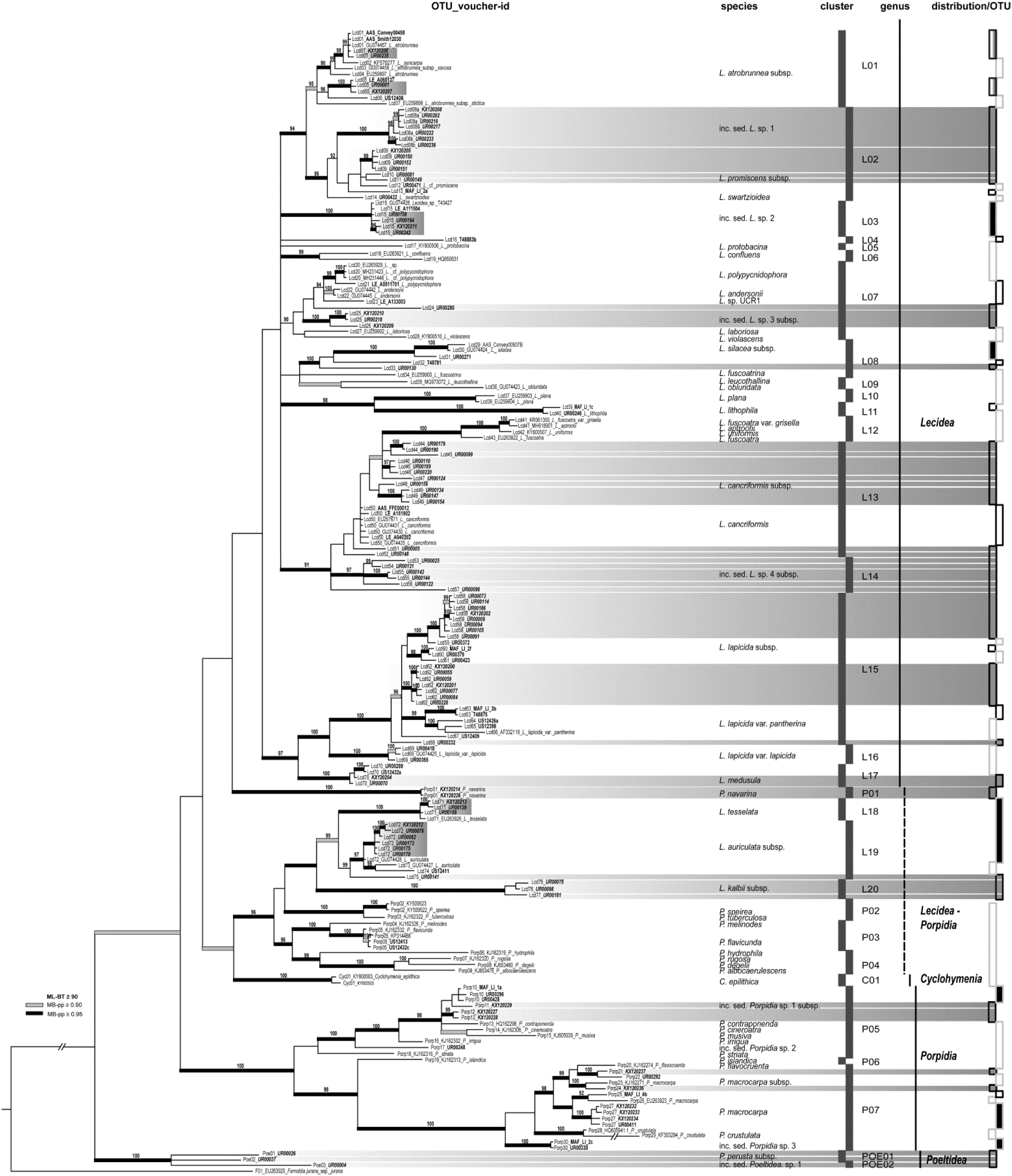
Phylogeny including all available relevant taxonomically identified sequences of the genera ***Lecidea, Porpidia, Poeltidea*** and ***Cyclohymenia*** using the **marker ITS**. Voucher information is preceded by the OTU number. The bars beside the phylogeny show the affiliation to clusters, genera and OTU distribution. sSA accessions are highlighted in grey and those restricted to sSA on OTU-level, with a full-length bar. The bootstrap values with ≥ 95 support of ML analyses were directly mapped on the Bayesian tree with ≥ 0.90 (gray) and ≥ 0.95 (black) support posterior probability values (branches in bold).

In all cases the ML analyses (IQ-TREE web server; Trifinopoulos et al. 2016) of the six phylogenies recovered the same tree topology as the MrBayes analysis (Ronquist & Huelsenbeck 2003). Therefore, we only present here the Bayesian tree with support values of the ML analyses.

For the multi-marker trees no conflict was found with the single marker trees, thus they are not shown. OTU scores with a sequence similarity higher than 97.5 % were calculated for groups with at least three accessions and are summarized in Table S5.

The assignment of uncertain accessions to mycobiont-species or subspecies was also based on morphological data, but these data are not included here.

### Lecidea/Porpidida/Poeltidea - ITS (Fig. 3)

This molecular phylogeny includes all relevant taxonomically identified sequences of the genera *Lecidea*, *Porpidia*, *Poeltidea* and *Cyclohymenia* provided by the project framework of the first author or downloaded from the NCBI database (GenBank) to bring the newly generated sequences of this study into a global context. The final data matrix for the phylogeny comprised 192 single sequences of the marker nrITS with a length of 593 characters and includes sequences of specimens of the genera *Lecidea* (144) *Porpidia* (42), *Poeltidea* (3), *Cylcohymenia* (2) and was rooted with *Farnoldia jurana* subsp. *jurana* as outgroup.

The phylogenetically delimited groups revealed were assigned to OTU-, species-, cluster- and genus level. All three analyses (distance-, model based and bGMYC) showed similar topologies, but at different levels. The groups gained with distance approach (OTUs) were used to show the intraspecific and/or the cryptic speciation; model based approaches were used for the assignment at species level and the clusters based on the bGMYC were used for grouping of closely related species or dividing highly heterogeneous species.

Altogether, 110 (*Lecidea*: 77; *Porpidia*: 31; *Cyclohymenia*: 1; *Poeltidea*: 3) OTUs were classified. These OTUs were assigned to species, including groups of subspecies and clusters (*Lecidea*: 20; *Porpidia*: 7; *Cyclohymenia*: 1; *Poeltidea*: 2).

The backbone of this phylogeny is not supported, but at least four main groups can be recognized. The first group, with low support, is formed solely by species of the genus *Lecidea* s. str. However, it can be considered a consistent group, because it is strongly supported by the three-marker phylogeny (Fig. S1). The southern hemisphere lineage (*P. navarina*) is situated outside the *Lecidea* group and next to the second group, which is an intermixed group of *Lecidea* and *Porpidia* species with two weakly supported accessions of the genus *Cyclohymenia* at the base. The third group is solely formed by species of the genus *Porpidia* and the fourth by the southern hemisphere genus *Poeltidea*.

The *Lecidea* group forms 17 clusters. The well supported, heterogeneous and cosmopolitan species cluster L01 (*L. atrobrunnea* subsp.) is sister to the well supported and diverse cluster L02 (inc. sed. *L.* sp. 1/endemic to sSA, *L. promiscens*/cosmopolitan and *L. swartzioidea*/northern hemisphere). Cluster L03 (inc. sed. *L.* sp. 2) forms a highly supported and very homogenous clade, comprising specimens from Antarctica, the Arctic and sSA. An unidentified specimen from continental Antarctica (T8883b) and an Arctic specimen of *L. protabacina* form clusters L04 and L05.

Cluster L06 includes *L. confluens* and an accession from Genbank (*Porpidia speirea*; Schmull et al. 2011). The placement of this latter accession in the *Lecidea* s. str. group could be caused by an incorrect species assignment (*Lecidea confluens* and *P. speirea* are morphologically quite similar and mainly distinguished by the ascus-type), or an uncertain species concept. The latter seems to not be the case, because two accessions of *P. speirea* from China were included in the phylogeny and are part of cluster P02, which agrees with the findings of Buschbom and Mueller (2004), who described this species as closely related to *P. tuberculosa*, which was confirmed in this phylogeny. Cluster L07 includes species related to *L. laboriosa* (northern hemisphere), such as the Antarctic and Chinese species *L. polypycnidophora* the bipolar *L. andersonii* and the Antarctic *L*. sp. UCR1. The Californian specimen of *Lecidea violascens* and an undetermined specimen from the Austrian Alps (*L.* sp. UR00280) are also part of this cluster.

Several well distinguished species form strongly supported clusters, such as the variable accessions from the northern hemisphere and maritime Antarctica of *L. silacea* subsp. together with an unidentified specimen form sSA (L. sp. UR00130) and *L. fuscoatrina* (cluster L08), followed by the northern hemisphere species *L. leucothallina* and *L. obluridata* (cluster L09). *Lecidea plana* (northern hemisphere; cluster L10) and *L. lithophila* (northern hemisphere, maritime Antarctica; cluster L11) form a highly supported clade.

Two recently newly described species, *Lecidea aptrootii* (Khan et al. 2018) and *L. uniformis* (McCune et al. 2017) are molecularly very closely related to the accession of *L. fuscoatra* var. *grisella* (Zhao et al. 2016), a taxon which was recognized by Aptroot & Van Herk (2007) at the species level as *L. grisella.* Together with *L. fuscoatra* they form the strongly supported cluster L12. This cluster is sister to cluster L13, which includes the Antarctic and, so far, endemic species *L. cancriformis* and a closely related, but heterogeneous subspecies group (*L. cancriformis* subsp.) from sSA. Another new and endemic species from sSA (inc. sed. *L.* sp. 4), forms the highly supported cluster L14.

Cluster L15 is formed by the very common, heterogeneous and cosmopolitan species group *L. lapicida* subsp. (including the cosmopolitan *L. lapicida* var. *pantherina*) and is sister to the northern hemisphere cluster L16, which consists of *L. lapicida* var. *lapicida.* These subspecies show no morphological differentiation and are separated only by their different chemotypes, but have according to Hertel (1995) different ecological requirements. The now cosmopolitan species *L. medusula,* previously only reported for the southern hemisphere (Fryday & Hertel 2014) forms cluster L17 with specimen from sSA, the Arctic and the Austrian Alps at the base of this clade. The species included in this group are part all referable to *Lecidea* s. str. (Hertel 1984).

The second group, which contains species of the genera *Lecidea* s. str. and *Porpidia*, is highly supported with the newly described species/genus *Cyclohymenia epilthica* (cluster C01) basal to the group, although with weak support. It is formed by two well-known and cosmopolitan species *L. tessellata* and *L. auriculata* (cluster L18 & L19) and the southern hemisphere *L. kalbii* (cluster 20) as well as a well-supported, northern hemisphere group of *Porpidia* species including *P. speirea, P. tuberculosa, P. melinodes* and *P. flavicunda* (cluster P02, P03) and other closely related species: *P. hydrophila, P. rugosa, P. degelii* and *P. albocaerulescens* (cluster P04).

However, species of the genus *Porpidia* are clearly distinguishable morphologically from species of *Lecidea*, by their ascus type and additionally by the associated green microalgae of the genus *Asterochloris* (Wirth et al. 2013; Ruprecht et al. 2016).

The third group is formed by three clusters of *Porpidia* species and is divided into two well supported main subgroups. One is formed by cluster P05 including the cosmopolitan species inc. sed. *P.* sp. 1 subsp. together with *P. cinereoatra, P. contraponenda, P. musiva, P. irrigua,* inc. sed. *P.* sp. 2 and *P. striata. Porpidia islandica* forms the highly supported cluster P06. The other strongly supported subgroup is formed by the heterogeneous and cosmopolitan cluster P07, including *P. flavocruenta, P. macrocarpa* subsp. and *P. macrocarpa*, the quite differing accessions from Turkey and China that were identified as *P. crustulata* and the inc. sed. *P.* sp. 3.

The genus *Poeltidea,* which occurs only in the southern hemisphere, forms clade IV and consists of two clusters, POE01 (*P. perusta*) and POE02 (inc. sed. *P*. sp. 1).

### Note

The family *Porpidaceae* was included in the synonymy to Lecideaceae (Lecideales) by Lumbsch & Huhndorf (2010), which is still the case in the current issue of Outline of *Ascomycota*: 2017 (Wijayawardene et al. 2018). The investigated genera *Lecidea* s. str. (Hertel 1984), *Porpidia*, *Poeltidea* as well as *Cyclohymenia* and *Farnoldia* are all assigned to this family, although *Farnoldia* appears to occupy a peripheral position.

### Lecidea/Porpidida/Poeltidea - ITS/mtSSU/RPB1 (Fig. S1)

The final data matrix of this phylogeny contains 204 concatenated sequences of the markers ITS, mtSSU and RPB1 with a length of 2115 characters and includes sequences of specimens of the genera *Lecidea*, *Porpidia* and *Poeltidea* and was rooted with *Farnoldia jurana* subsp. *jurana* as outgroup. Specimens from the project framework of the first author (Antarctica, Arctic, Austria) with the same three markers were added.

All available sequences of this study were included to show the abundance of the different specimens in the studied areas of sSA. The phylogenetically delimited groups revealed were assigned to OTU-, species-, and genus level. Additionally, the distribution of the OTUs is shown beside the genus information.

This multi-marker phylogeny is not fully comparable to the overall single marker (ITS) phylogeny of *Lecidea/Porpidida/Poeltidea* because of the limited availability of sequences of the chosen markers in GenBank. The topology is, in most cases, similar and it forms the same four groups, but in many cases, they show greater support. The *Lecidea* group, at least, is strongly supported in this phylogeny. Because of the limited availability of *Porpidia* sequences in GenBank, the topology of this group is slightly different with *P. navarina* clustering together with *P. cinereoatra* and *P. macrocarpa*, but with low support.

Finally, the genus *Poeltidea* with two species is still at the base of this phylogeny.

The OTUs obtained from the ITS sequences are still very well supported. Many clades show clearly a local differentiation at OTU level and are endemic to sSA. The two largest clades (OTUs Lcd58 and Lcd62) are part of the cosmopolitan *L. lapicida* cluster and are the most abundant accessions. These are followed by *Porpidia navarina* and inc. sed. *Lecidea* sp. 1 (both species endemic to sSA), *L. promiscens* (cosmopolitan species group) and other smaller groups.

### *Lecidella* - ITS (Fig. 4)

To bring the new accessions of *Lecidella* from sSA into a global context, the species concept and most of the published accessions of Zhao et al. (2015) were used. Additionally, to this dataset several sequences from Genbank and from the project framework of the first author were added. The final data matrix of this phylogeny contains 76 sequences of the marker ITS with a length of 538 characters and was rooted with species of the genera *Carbonea, Lecanora* and *Rhizoplaca* to obtain well-defined units in the genus *Lecidella*.

The backbone of the phylogeny is unresolved, but four strongly supported main clades are formed; three (*L. enteroleucella, L. stigmatea* and *L. elaeochroma*) are the same as those identified by Zhao *et al*. (2015) plus a fourth clade (*L.* sp. nova) with two species only occurring in Antarctica and sSA. All available sequences for this group were included to show the abundance of the different accessions. The phylogenetically delimited groups revealed were assigned to OTU-, clade-, and genus level. Additionally, the global distribution of the locally differentiated OTUs is shown beside the genus information.

*Lecidella enteroleucella* is still the only member of the first clade.

The second clade (*L. stigmatea)* is completely unresolved. *Lecidella patavina* and *L. stigmatea* are intermixed and not assignable. These two species differ most noticeably in that the hymenium of *L. patavina* is inspersed with oil droplets, whereas that of *L. stigmatea* is not. That the accessions of these two species are intermixed perhaps indicates that either only one species is involved or that the defining character has been interpreted inconsistently. *Lecidella greenii* and *L. siplei* form well supported lineages and the 12 sequences of the species in this study show a clear local differentiation at OTU level.

A third, new and strongly supported, clade (*L.* sp. nova) shows two new species from sSA (*L.* sp. 1) and continental Antarctica (*L.* sp. 2).

The fourth clade (L. elaeochroma) shows a few separated and well supported species (L. tumidula, L. meiococca, L. wulfenii, L. flavosorediata and L. sp. 3, which is endemic to sSA). Lecidella elaeochroma, L. euphorea, L. carpathica, L. elaeochromoides and L. effugiens are not assignable because of mingling in different highly supported lineages. None of the investigated specimens were morphologically similar to the southern hemisphere species Lecidella sublapicida (Knoph & Leuckert 1994).

### Asterochloris - ITS (Fig 5)

All the *Porpidia* species in this study are not only associated with *Trebouxia* as photobiont, but also with a green microalgae of the genus *Asterochloris.* The accessions obtained from sSA were brought into a global context by adding all relevant taxonomically identified sequences from GenBank and from the project framework of the first author. The final data matrix of this phylogeny contains 73 sequences of the marker ITS with a length of 519 characters.

The phylogeny was rooted midpoint and divided into two main clades (genera), *Asterochloris* and *Vulcanochloris*. The accessions from sSA occur only in the *Asterochloris* clade. The backbone of this clade is unresolved. Again, all available sequences for this group are included to show the abundance of the different accessions. The phylogenetically delimited groups revealed were assigned to OTU-, clade-, and genus level. Additionally, the global distribution of the locally differentiated OTUs is shown beside the genus information.

The topology of the species from GenBank show a similar pattern to that already described in Ruprecht et al. (2014).

Only one accession from sSA clusters together with the cosmopolitan species *A. woessiae*. A highly supported and homogeneous clade is formed by 19 specimens (*A.* sp. URa18) only occurring on Isla Navarino and one accession from the other side of the Beagle channel (Tierra del Fuego) in the southernmost part of sSA. Four other accessions (*A.* sp. URa19, *A.* sp. URa20, *A.* sp. UR00027 and *A*. sp. UR00123) are placed in the main group with low support.

### *Trebouxia* - ITS (Fig. 6)

To bring the new accessions of *Trebouxia* from sSA into a global context, the species/OTU concept and the dataset reduced to one accession of each OTU of Leavitt et al. (2015) was used. Additionally, several new sequences from GenBank and from the project framework of the first author were added to this dataset. The final data matrix of this phylogeny contains 159 sequences of the marker ITS with a length of 803 characters and was midpoint-rooted.

The phylogenetically delimited groups revealed were assigned to OTU- and clade level as described in Leavitt et al. (2015).

The backbone of the phylogeny is unresolved, but five strongly supported main clades were formed; four clades (I, A, G and S) correspond to those of Leavitt et al. (2015) plus a fifth clade (N) with three specimens only occurring in sSA on Isla Navarino and on the north side of the Beagle channel at Tierra del Fuego.

Three different groups were formed by accessions assigned to Clade I. Because of a sequence similarity below the threshold of 97.5 % the subOTU I01i was divided into two subunits (I01i, I01j) with a cosmopolitan and southern Polar distribution, plus an independent OTU I17 solely occurring in sSA. Due to the addition of the diverse accessions from sSA to the existing OTU I01, which already has 10 subOTUs (I01, I01a – i) described by Leavitt et al. (2015), most of them were transferred into distinct OTUs. However, these have not been renamed here, as this is an open system and a regrouping with new accessions is expected in the future.

Altogether, clade A includes 39 OTUs with the specimens of this study forming part of two cosmopolitan (A02, A04) and four locally differentiated (A36 - A39) OTUs. A04 has a sequences similarity of 96.7% and was subdivided in two subunits (A04a, A04b).

Fifty percent of the sSA accessions of this marker are contained in the cosmopolitan OTUs S02 and S07 of clade S. S02 was subdivided into four subunits with the accessions from sSA only being assigned to S02 together with northern hemisphere specimens. S02b is solely formed by specimen from continental Antarctica, a similar finding was described in Ruprecht et al. (2012a; *T. jamesii* subsp. is equivalent to S02b in this study). Another strongly supported OTU (S16) consists only of accessions from sSA.

The new and highly supported clade N appeared with two haplotypes (three accessions), all of which are located at the southernmost part of sSA.

No accession from this study is part of clade G.

### *Trebouxia* - ITS/psbJ-L/COX2 (Fig. S2)

The data matrix of this phylogeny contains 217 concatenated sequences of the markers ITS, psbJ-L and COX2 with a length of 1693 characters. Only sSA specimens are included to demonstrate the intraspecific differentiation. This dataset was calibrated with four cultured *Trebouxia* specimens (Table S2d, 2×S02/Antarctica, A02/Antarctica and A12/Sweden). Interestingly, the psbJ-L sequences of A02 of the cultured specimen from Antarctica are different to the North American specimen from Leavitt et al. (2015).

The dataset still shows the same number of OTUs as the ITS phylogeny (Fig. 6) and the grouping is the same. Clade A shows nine well supported groups at species level (A02, A12, A38, A39, A04a, A04b, A36, A37) with A36 and A04b being closely related. Interestingly, the subOTUs A04a and A04b are more separated with the marker psbJ-L than in the ITS phylogeny.

Clade I is divided into two subOTUs (I01i, I01j) and one newly developed distinct OTU (I17) because of its heterogeneous structure. No COX2 sequences were included, because they were not assignable.

More than half of the sequences are included in the homogenous and cosmopolitan OTUs S02 and a smaller part in S07. These groups are closely related and share similar COX2 sequences. The OTU S16, which occurs only in sSA, is clearly separated at species level from S02 and S07. The new clade N is still strongly supported in this dataset, but basal to clade S.

### Distribution of species and OTUs: globally distributed vs. restricted to sSA and/or southern Polar regions

Altogether, 185 mycobiont specimens forming 54 OTUs assigned to 24 species of the genera *Lecidea, Porpidia, Poeltidea* and *Lecidella* were identified.

Four species of the genus *Lecidea* (inc. sed. *L.* sp. 2, *L. auriculata, L. medusula, L. tessellata*) and *Porpidia macrocarpa* are globally distributed at species and OTU level.

By far the most abundant accessions are formed by locally differentiated OTUs (Lcd58, Lcd62; 48 accessions) and belong to the cosmopolitan cluster *Lecidea lapicida* (L15), with the next most frequent groups being *Lecidella stigmatea* (13) and *Lecidea promiscens* (9). Finally, altogether 13 species or single sequences are so far only described and/or known from sSA or the southern Polar regions (Table 1, Figs. 3, S1 & 4).

**Table 1:** Mycobiont distribution of species and OTUs. Species and OTUs with different distributions (globally/sSA) are marked in bold.

| species - distribution |  | n | OTU - distribution |  |
| --- | --- | --- | --- | --- |
| inc. sed. *Lecidea sp. 2* | globally | 8 | LP_Lcd15 | globally |
| *Lecidea auriculata* | globally | 13 | LP_Lcd72 | globally |
| *Lecidea medusula* | globally | 3 | LP_Lcd70 | globally |
| *Lecidea tessellata* | globally | 3 | LP_Lcd71 | globally |
| *Porpidia macrocarpa* | globally | 4 | LP_Porp27 | globally |
| **Subtotal** | **5** | **31** | **5** |  |
| *Lecidea atrobrunnea* subsp. | globally | **5** | LP_Lcd01, 05 | **southern polar** |
| *Lecidea lapicida* subsp. | globally | **48** | LP_Lcd58, 62, 68 | **sSA** |
| *Lecidea promiscens* subsp. | globally | **9** | LP_Lcd09 - 11 | **sSA** |
| *Lecidella stigmata* | globally | **13** | LL_ST07, 08, 09, 15 - 20 | **sSA** |
| inc. sed. *Porpidia sp. 1* subsp. | globally | **5** | LP_Porp11, 12 | **sSA** |
| *Porpidia macrocarpa* subsp. | globally | **2** | LP_Porp21, 24 | **sSA** |
| **Subtotal** | **6** | **81** | **21** |  |
| inc. sed. *Lecidea* sp. 1 | sSA | 8 | LP_Lcd08a, 08b | sSA |
| inc. sed. *Lecidea* sp. 3 | sSA | 4 | LP_Lcd25, 26 | sSA |
| inc. sed. *Lecidea* sp. 4 | sSA | 8 | LP_Lcd53 - 56 | sSA |
| inc. sed. *Lecidella* sp. 1 | sSA | 3 | LL_N01 | sSA |
| inc. sed. *Lecidella* sp. 3 | sSA | 2 | LL_EL25 | sSA |
| inc. sed. *Poeltidea* sp.1 | sSA | 3 | LP_Poe03 | sSA |
| *Lecidea cancriformis* subsp. | sSA | 19 | LP_Lcd44 – 49, 51, 52 | sSA |
| *Lecidea kalbii* | sSA | 4 | LP_Lcd76, 77 | sSA |
| *Lecidea* sp. UR00096 | sSA | 1 | LP_Lcd57 | sSA |
| *Lecidea* sp. UR00130 | sSA | 1 | LP_Lcd33 | sSA |
| *Lecidea* sp. UR00141 | sSA | 1 | LP_Lcd75 | sSA |
| *Porpidia navarina* | sSA | 13 | LP_Porp01 | sSA |
| *Poeltidea perusta* | sSA | 5 | LP_Poe01, 02 | sSA |
| **Subtotal** | **13** | **73** |  | **28** |
| **Total:** | **24** | **185** |  | **54** |

**Figure 4:**
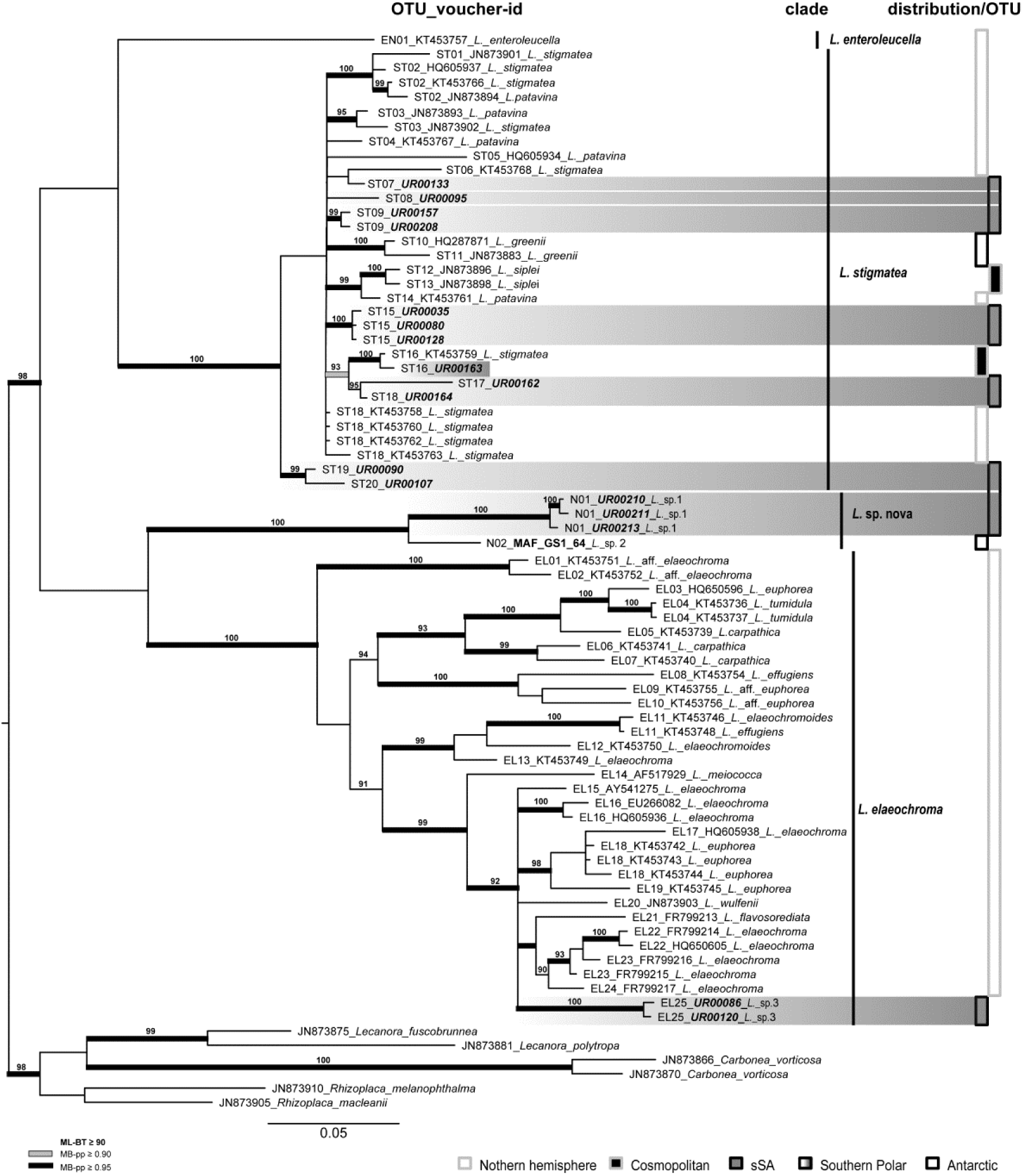
Phylogeny of the genus ***Lecidella*** with accessions from sSA integrated in the species concept and most of the published accessions (Zhao et al. 2015) using the **marker ITS**. Voucher information is preceded by the OTU number. The bars beside the phylogeny show the affiliation to clades and distribution. sSA accessions are highlighted in grey and those restricted to sSA on OTU-level, with a full-length bar. The bootstrap values with ≥ 95 support of ML analyses were directly mapped on the Bayesian tree with ≥ 0.90 (gray) and ≥ 0.95 (black)support posterior probability values (branches in bold).

The photobionts comprise 199 accessions that are assigned to 19 OTUs. Most of the algal specimens (125) belong to globally distributed taxa, especially Tr_S02a with 84 accessions followed by Tr_S07, Tr_A02, Tr_A12, Tr_A04a and a single accession of *Asterochloris woessiae* from a lower elevation area close to Esquel, Argentina. The very heterogeneous OTU Tr_I0i was divided into two sub-OTUs: I01i with a global distribution and I01j that occurs in Antarctica and sSA. OTU I17 occurs solely in sSA. A surprisingly high number of 82 accessions (*Asterochloris* and *Trebouxia*) form 13 clearly separated OTUs and were, so far, only found in sSA (Table 2, Figs. 5, 6 & S2).

**Table 2:** Photobiont distribution of species and OTUs

| species | n | OTU - distribution |  |
| --- | --- | --- | --- |
| *A. woessiae* | 1 | Ast13 | globally |
| *Trebouxia* sp. | 7 | Tr_A02 | globally |
| *Trebouxia* sp. | 5 | Tr_A04a | globally |
| *Trebouxia* sp. | 6 | Tr_A12 | globally |
| *Trebouxia* sp. | 5 | Tr_I01i | globally |
| *Trebouxia* sp. | 84 | Tr_S02 | globally |
| *Trebouxia* sp. | 17 | Tr_S07 | globally |
| **Subtotal** | **125** | **7** |  |
| *A. sp.* UR00027 | 1 | Ast22 | sSA |
| *A. sp.* UR00123 | 1 | Ast21 | sSA |
| *A. sp.* URa18 | 19 | Ast24 | sSA |
| *A. sp.* URa19 | 1 | Ast23 | sSA |
| *Trebouxia* sp. | 6 | Tr_A04b | sSA |
| *Trebouxia* sp. | 1 | Tr_A36 | sSA |
| *Trebouxia* sp. | 2 | Tr_A37 | sSA |
| *Trebouxia* sp. | 4 | Tr_A38 | sSA |
| *Trebouxia* sp. | 5 | Tr_A39 | sSA |
| *Trebouxia* sp. | 20 | Tr_I01j | southern Polar |
| *Trebouxia* sp. | 11 | Tr_I17 | sSA |
| *Trebouxia* sp. | 3 | Tr_S16 | sSA |
| **Subtotal** | **74** | **12** |  |
| **Total** | **199** | **19** |  |

Summarizing, for the mycobiont the percentage distribution of accessions at OTU level shows a high rate of local differentiation and endemism for the sSA specimens vs those that are globally distributed (**83**:17). In particular, Parque Nacional Torres del Paine and Morro Chico with 100% each, and the southernmost sampling point, Isla Navarino with 90% have the highest amount of specialized accessions. In contrast, the photobiont OTUs show a higher rate of globally distributed accessions (32:**68**). However, both symbionts show no significant specialization along the latitudinal gradient in southern South America (Table 3).

**Table 3:** Summary of investigation sites with elevation, climate variables BIO1 (annual mean temperature) and BIO 12 (annual precipitation), using CHELSA (Karger et al. 2017), and information about locally differentiated OTUs and/or endemic species.

| location | latitude (°S) | altitude m a.s.l. | climate variables |  | locally diff. OTUs and/or endemic species to sSA (yes/no - proportional) |  |  |  |
| --- | --- | --- | --- | --- | --- | --- | --- | --- |
|  |  |  | Bio 1 (°C) | Bio 12 (mm) | mycobiont |  | photobiont |  |
|  |  |  |  |  | n | yes no | n | yes no |
| Bariloche (AR) | 41.2 | 2001 to 2061 | 1.7 | 1642 | 18 | 56 44 | 18 | 17 83 |
| Esquel (AR) | 42.8 | 1964 to 2122 | 1.2 to 1.4 | 1140 to 1180 | 26 | 69 31 | 26 | 23 77 |
| Tecka-Corcovado (AR) | 43.5 | 797 | 7.8 | 685 | 9 | 89 11 | 9 | 56 44 |
| PN Los Glaciares (AR) | 49.2 to 50.5 | 558 to 1481 | 0.2 to 4.1 | 322 to 512 | 43 | 81 19 | 40 | 32 68 |
| PN Torres del Paine (CL) | 51.1 to 51.6 | 118 to 324 | 3.9 to 6.5 | 545 to 818 | 12 | 100 | 13 | 38 62 |
| Morro Chico (CL) | 52 | 208 | 6.4 | 341 | 10 | 100 | 9 | 44 56 |
| Ushuaia (AR) | 54.7 to 54.8 | 407 to 794 | 1.6 to 4.0 | 489 to 756 | 34 | 76 24 | 35 | 11 89 |
| Isla Navarino (CL) | 55 | 607 to 741 | 1.9 to 3.3 | 628 to 739 | 33 | 90 10 | 52 | 38 62 |
| **Total (mean)** |  |  |  |  | **185** | **83 17** | **202** | **32 68** |

**Figure 5:**
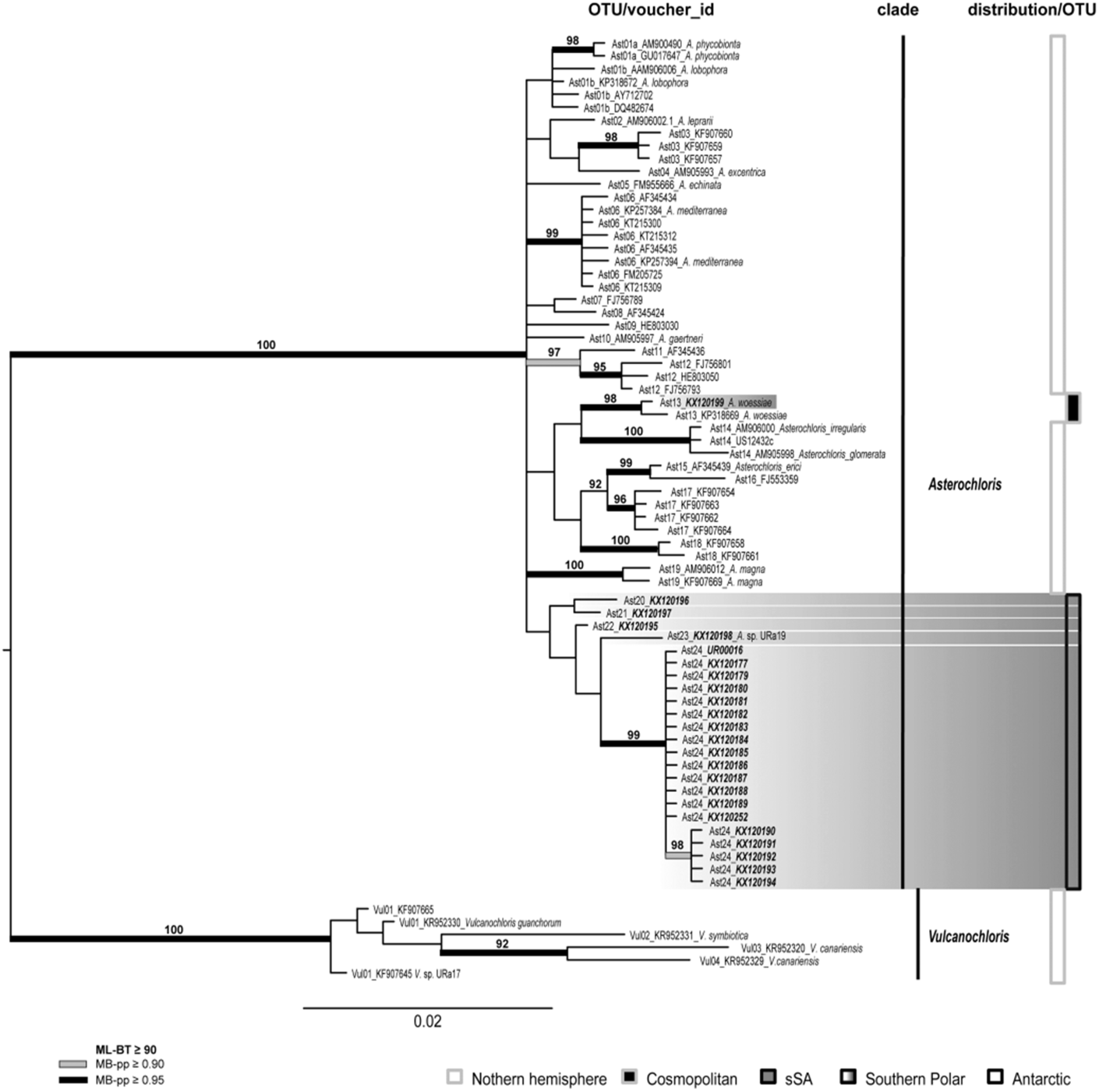
Phylogeny of the genus ***Asterochloris*** including all available relevant taxonomically identified sequences using the **marker ITS**. Voucher information is preceded by the OTU number. The bars beside the phylogeny show the affiliation to genera and distribution. sSA accessions are highlighted in grey and those restricted to sSA on OTU-level, with a full-length bar. The bootstrap values with ≥ 95 support of ML analyses were directly mapped on the Bayesian tree with ≥ 0.90 (gray) and ≥ 0.95 (black) support posterior probability values (branches in bold).

**Figure 6:**
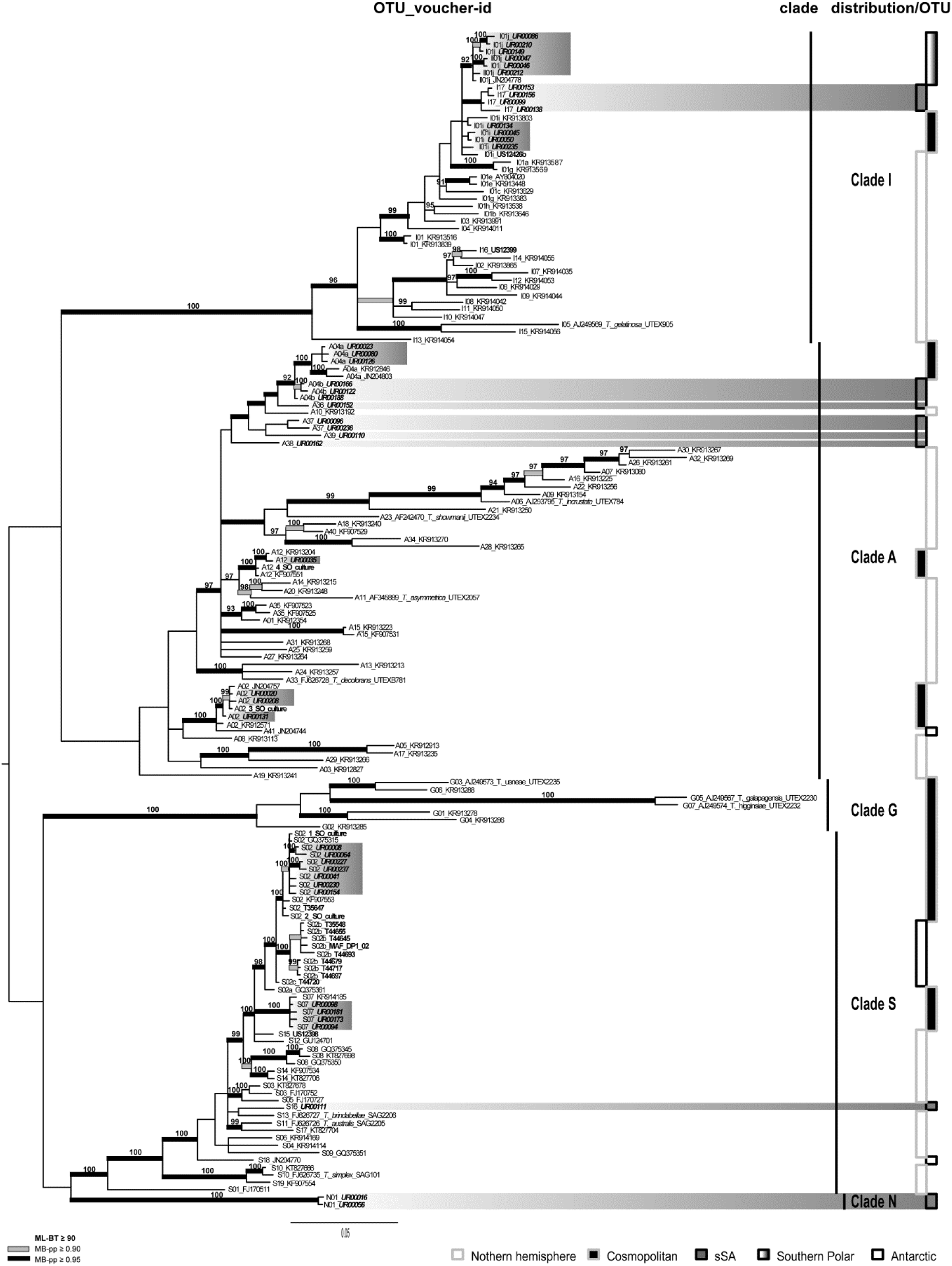
Phylogeny of the genus ***Trebouxia*** with accessions from SSA integrated in to the species concept and most of the published accessions of Leavitt et al. 2015 using the **marker ITS**. Voucher information is preceded by the OTU number. The bars beside the phylogeny show the affiliation to the clades. sSA accessions are highlighted in grey and those restricted to sSA on OTU-level, with a full-length bar. The bootstrap values with ≥ 95 support of ML analyses were directly mapped on the Bayesian tree with ≥ 0.90 (gray) and ≥ 0.95 (black) support posterior probability values (branches in bold).

## Discussion

For the four re-evaluated groups *Lecidea/Porpidia/Poeltidea*, *Lecidella* (mycobiont), *Asterochloris* and *Trebouxia* (photobiont), the geographical isolated southern end of the South American continent supports a high degree of locally differentiated subclades (OTUs) in globally distributed species as well as endemic lineages at the species and genus level. This was, to some extent, unexpected for these mostly globally distributed genera and can partially be explained by the lack of sequence information for most of the southern hemisphere lecideoid mycobiont species (Fryday & Hertel 2014; Knoph & Leuckert 1994). This also applies to the photobionts because of the limited availability of molecular studies for the southern Polar Regions (e.g. Fernandez-Mendoza *et al*. 2011; Muggia *et al*. 2010; Ruprecht *et al*. 2012a).

However, local differentiation (cryptic speciation) is quite common in lichen forming fungi (Dal Grande et al. 2017; Kraichak et al. 2015; Leavitt et al. 2011; Lumbsch and Leavitt 2011), but has rarely been described for the most common and widespread lichen photobiont taxa *Trebouxia* (Fernandez-Mendoza et al. 2011; Leavitt et al. 2015; Ruprecht et al. 2012a) and *Asterochloris* (Skaloud et al. 2015).

In particular, the lichen cluster (L15, Fig. 3) *Lecidea lapicida*, which occurs in polar and high mountainous regions worldwide (Hafellner and Türk 2016; Hertel 1984; Hertel and Andreev 2003), shows locally differentiated accessions at the OTU level that are endemic to sSA. Austrian and Antarctic accessions are closely related, but they are clearly distinct from the two main OTUs (Lcd58, Lcd62) that comprise 48 of the 185 sSA specimens. The heterogeneous clades of *Porpidia macrocarpa* subsp., which include specimens from Antarctica and the Austrian Alps, show a similar pattern. *Lecidea atrobrunnea* subsp. is the only exception, forming two southern polar OTUs with accessions from both Antarctica and sSA (Fig. 3). The same pattern is also known for *Usnea aurantiacoatra* (Laguna Defior 2016) and *Cetraria aculeata* (Fernandez-Mendoza et al. 2011). A different example is *Lecidea cancriformis*, which was, so far, described as endemic to Antarctica and is one of the dominant crustose lichens in the most extreme areas of the continent (Castello 2003; Hertel 2007; Ruprecht et al. 2010). The Antarctic accessions belong to a single, well supported OTU (Lcd50, Fig. 3), but there are seven closely related OTUs occurring in sampling areas north of Tierra del Fuego. However, the whole cluster L13 including *L. cancriformis* subsp. remains confined to the southern Polar Regions. Several other species, as well as the genus *Poeltidea,* occur solely in the southern Polar Regions. *Lecidea medusula,* which was previously only investigated morphologically and thought to be endemic to the southern hemisphere (Hertel 2009), is shown to be a cosmopolitan species. However, in total, the cosmopolitan species/OTUs (e.g. *L. auriculata, L. tessellata*) are outnumbered by those that are endemic to sSA (Table 1).

The *Lecidella* phylogeny, based on the data of Zhao et al. (2015), reveals a new southern polar species-level clade (*L.* sp. nova) with accessions from sSA and continental Antarctica. The specimens are not morphologically assignable to the available species descriptions (e. g. Knoph & Leuckert 1994; Ruprecht et al. 2012b; Wirth et al. 2013). All the other sequences added to Zhao et al.’s phylogeny form well supported and distinguished OTUs endemic to sSA.

Interestingly, several bipolar (incl. high mountainous areas worldwide) and abundant mycobiont species occurring in continental and maritime Antarctica, for example *Lecidea andersonii, L. polypycnidophora* (Hertel 2007; Ruprecht et al. 2010, Ruprecht et al. 2016), *Lecidella siplei* (Ruprecht et al. 2012b) were not found in the sSA regions.

In contrast to the mycobiont, the photobiont shows the opposite pattern. In particular, the genus *Trebouxia* is known as globally widely distributed with often low diversification (Muggia et al. 2010; Ruprecht et al. 2012a). These findings are supported in this study with more than half of the accessions assigned to two cosmopolitan OTUs of the genus *Trebouxia* (S02, S07) plus some smaller groups (A02, A04a, A12; Leavitt et al. 2015; Figs. 6, S2, Table 2). The remaining *Trebouxia* and *Asterochloris* accessions form highly diverse and locally differentiated and/or endemic groups, including a new clade from Isla Navarino (*Trebouxia* clade N), which was surprising at this unprecedented scale. The contrasting distribution behavior of the cosmopolitan photobionts could be caused mainly by their wide choice of mycobiont partners (Kroken and Taylor 2000) allowing them access to the different distribution strategies of the various lichens.

The most southern sampling area at Isla Navarino shows, for the mycobiont, not only locally restricted OTUs, but also strongly supported endemic species (e.g. *Porpidia navarina*; Ruprecht et al. 2016), which is also the case for *Asterochloris* (*A.* sp. URa18) and for *Trebouxia* (clade N). As this area was ice-free during the Last Glacial Maximum (Douglass et al. 2005) the likely reason can be explained by the concept of glacial refugia where the cold and glacial phases were the drivers of population divergences and (cryptic) speciation after transition from the northern to the southern hemisphere (Fernandez-Mendoza and Printzen 2013; Paula and Leonardo 2006; Stewart et al. 2010). Furthermore, the two southernmost areas on the South American continent sampled (Parque Nacional Torres del Paine and Morro Chico), which both have 100% locally differentiated OTUs and endemic species for the mycobiont (Table 3), are influenced by the violent westerly gales caused by the split of the Humboldt current to the north and the Antarctic circumpolar current (ACC) to the south (Silva et al. 2009). This further leads to the assumption that mycobiont dispersal is limited through this asymmetrical wind system, which is caused by the undertow of the ACC, driven by westerly winds over the circumpolar streamlines (Allison et al. 2010). Moreover the flow speed of the ACC between the Last Glacial Maximum and the Holocene has remained almost unchanged (McCave et al. 2014). Nonetheless, several other strongly supported species groups that only occur in sSA in other areas, such as *Lecidella* sp. 1 at 2000 m a.s.l. close to Esquel (S42.8°) or endemic lineages in *Trebouxia* (S16, A38, A39), hint to further, and so far unknown, separation events at the remote and climatically extreme southern end of the American continent.

### Taxonomy

It is well known that the mycobiont genera *Lecidea* and *Porpidia* (Fig. 3) are not clearly separated and our phylogeny confirms that the species currently included in *Porpidia* do not form a monophyletic group (i.e. Buschbom & Mueller 2004; Schmull et al. 2011; Ruprecht et al. 2016; Fig. 3 & S1). Additionally, the newly described species/genus *Cyclohymenia epilithica* with perithecioid apothecia and an apparently *Porpidia* - type ascus (McCune et al. 2017) is situated among these two genera. In general, species of the genera *Lecidea* and *Porpidia* are morphologically differentiated by ascus-type (*Lecidea/Porpidia*), larger ascospores with the presence of a perispore in *Porpidia* and different genera of associated green microalgae as photobionts (*Trebouxia* sp. in *Lecidea* sp.; *Asterochloris* sp. and *Trebouxia* sp. in *Porpidia sp*.; Ruprecht et al. 2016). *Chlorella* sp. as described by Li et al. (2013) was not found in the sSA *Porpidia* species. Although, several new sequences were added for previously only morphologically described species e.g. *L. kalbii*, *L. promiscens*, *L. swartzioidea, L. lithophila*, *Poeltidea perusta*, and sequences for other *Porpidia* species obtained from Genbank, the re-evaluated phylogeny could not be resolved. Still, three species, morphologically assigned to *Lecidea* s. str. (*L. auriculata, L. tessellata, L. kalbii*; Fryday and Hertel 2014; Hertel 1984; Wirth et al. 2013) form, together with several *Porpidia* species, a highly supported, but intermixed group. Although clearly defined groups for *Porpidia* are easily recognized and a name at the genus level is available for at least one of them (*Haplocarpon*) it would be premature to formally recognize these groups as genera because of the uncertain systematic position of the type species of the genus, *Porpidia trullisata*, a rare species for which molecular data is not yet available. However, the three other groups are formed solely by species of *Lecidea* s. str., *Porpidia,* and *Poeltidea*, respectively. The unresolved and intermixed topology of several species in the two main clades of the *Lecidella* phylogeny (*L. stigmatea* and *L. elaeochroma*, Fig. 4) could not be improved with the additional specimens from sSA. Only an extended species sampling can help to unravel the inconsistent relationships in both these phylogenies (*Lecidea/Porpidida/Poeltidea*, Fig. 3; *Lecidella*, Fig. 4).

## Conclusions

The species-rich group of lecideoid lichens found widespread in Alpine and Polar Regions in southern South America comprises highly divergent OTUs of cosmopolitan species as well as several endemic species. Three factors may contribute to the observed differentiation and endemism: a) the geographical isolation of this southernmost landmass north of Antarctica, b) limited dispersal caused by the Antarctic circumpolar current system, and c) the presence of regional glacial refugia.

The diverging patterns of dispersal in the cosmopolitan lecideoid lichen group are still under researched. Gaining larger datasets along the assumed distribution routes of the highest mountain ranges (Garrido-Benavent & Perez-Ortega, 2017) and a consequent sampling for a better global coverage will help to understand colonization events and specialization in this, so far, quite overlooked group of crustose lichens.

## Acknowledgements

We are grateful to M. Affenzeller, H.P. Comes, M. Wagner (University of Salzburg), C. Printzen (Senckenberg, Frankfurt) and L. Sancho (Universidad Complutense Madrid) who gave us highly appreciated advice and support. G. Brunauer (University of Salzburg), A. Passo (University of Comahue), M. Andreev (Bot. Institute, St. Petersburg), British Antarctic Survey (Cambridge) and U. Søchting (University of Copenhagen) thankfully helped with collecting and providing the lichen specimens. Finally, we would like to thank Sieglinde Ott and Nadine Determeyer-Wiedmann (University of Düsseldorf) for providing several *Trebouxia*-cultures.

This study was financially supported by the Austrian Science Fund (FWF): P26638.

## Supplementary Information

**Figure S1:**
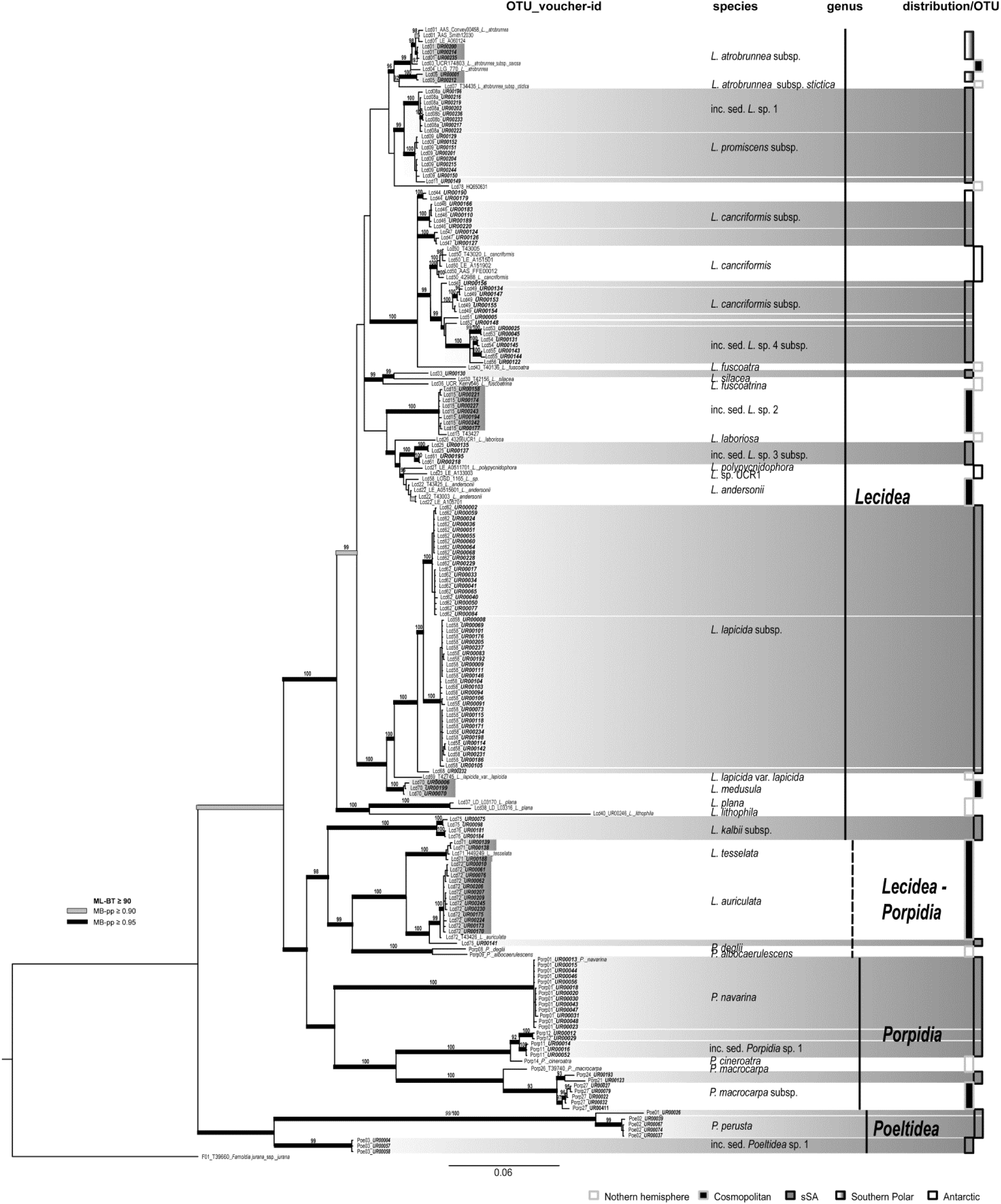
Phylogeny of concatenated **ITS, mtSSU** and **RPB1** sequences including all investigated specimens from sSA, and additional accessions from Antarctica and Europe of the genera ***Lecidea*, *Porpidia*** and ***Poeltidea***. Voucher information is preceded by the OTU number. The bars beside the phylogeny show the affiliation to genera and OTU distribution. sSA accessions are highlighted in grey and those restricted to sSA on OTU-level, with a full-length bar. The bootstrap values with ≥ 95 support of ML analyses were directly mapped on the Bayesian tree with ≥ 0.90 (gray) and ≥ 0.95 (black) support posterior probability values (branches in bold).

**Figure S2:**
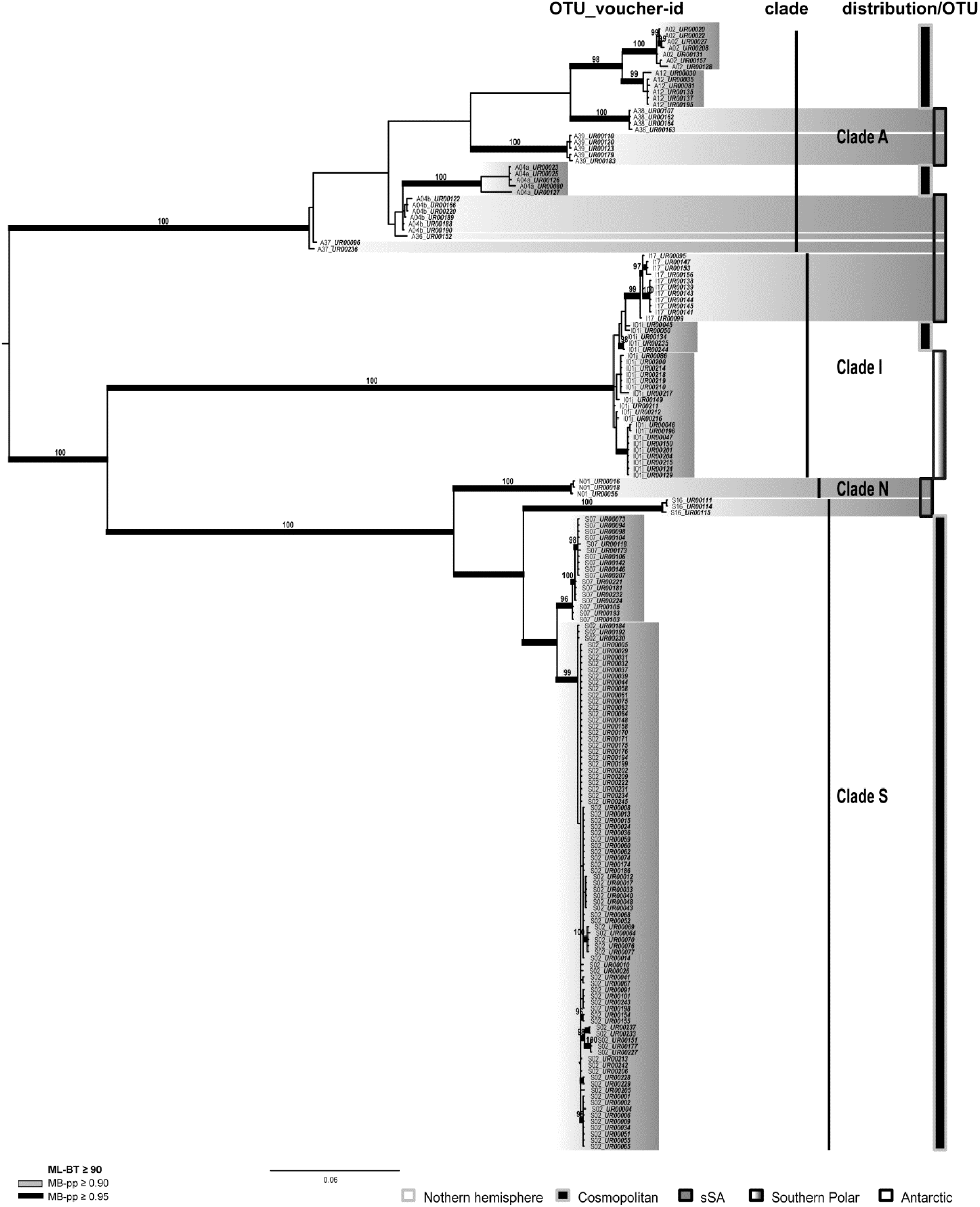
Phylogeny of concatenated **ITS, psbJ-L** and **COX2** sequences showing only the investigated specimens from sSA of the genus ***Trebouxia***. Voucher information is preceded by the OTU number. The bars beside the phylogeny show the affiliation to the clades and distribution. sSA accessions are highlighted in grey and those restricted to sSA on OTU-level, with a full-length bar. The bootstrap values with ≥ 95 support of ML analyses were directly mapped on the Bayesian tree with ≥ 0.90 (gray) and ≥ 0.95 (black)support posterior probability values (branches in bold).

**Table S1:**
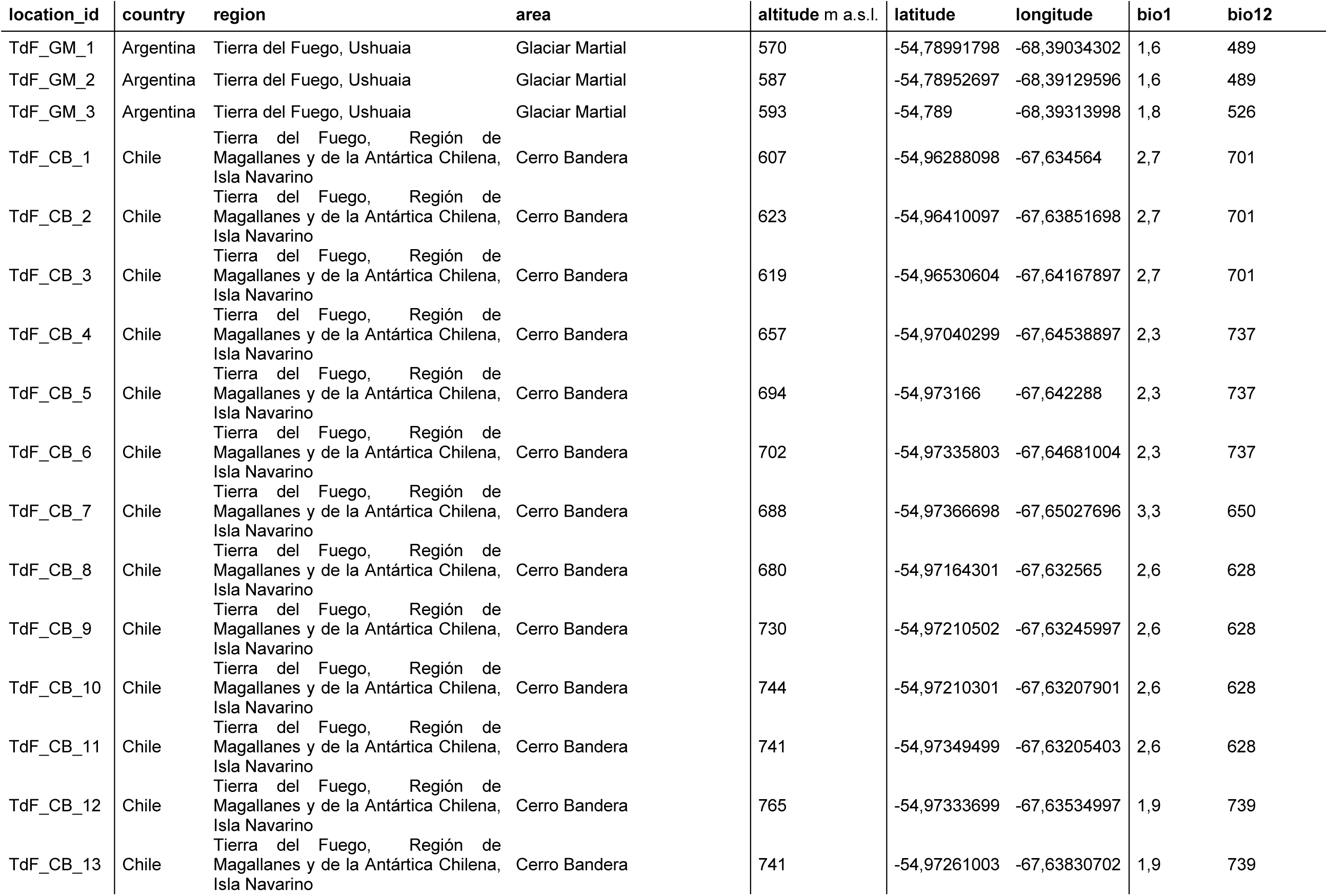
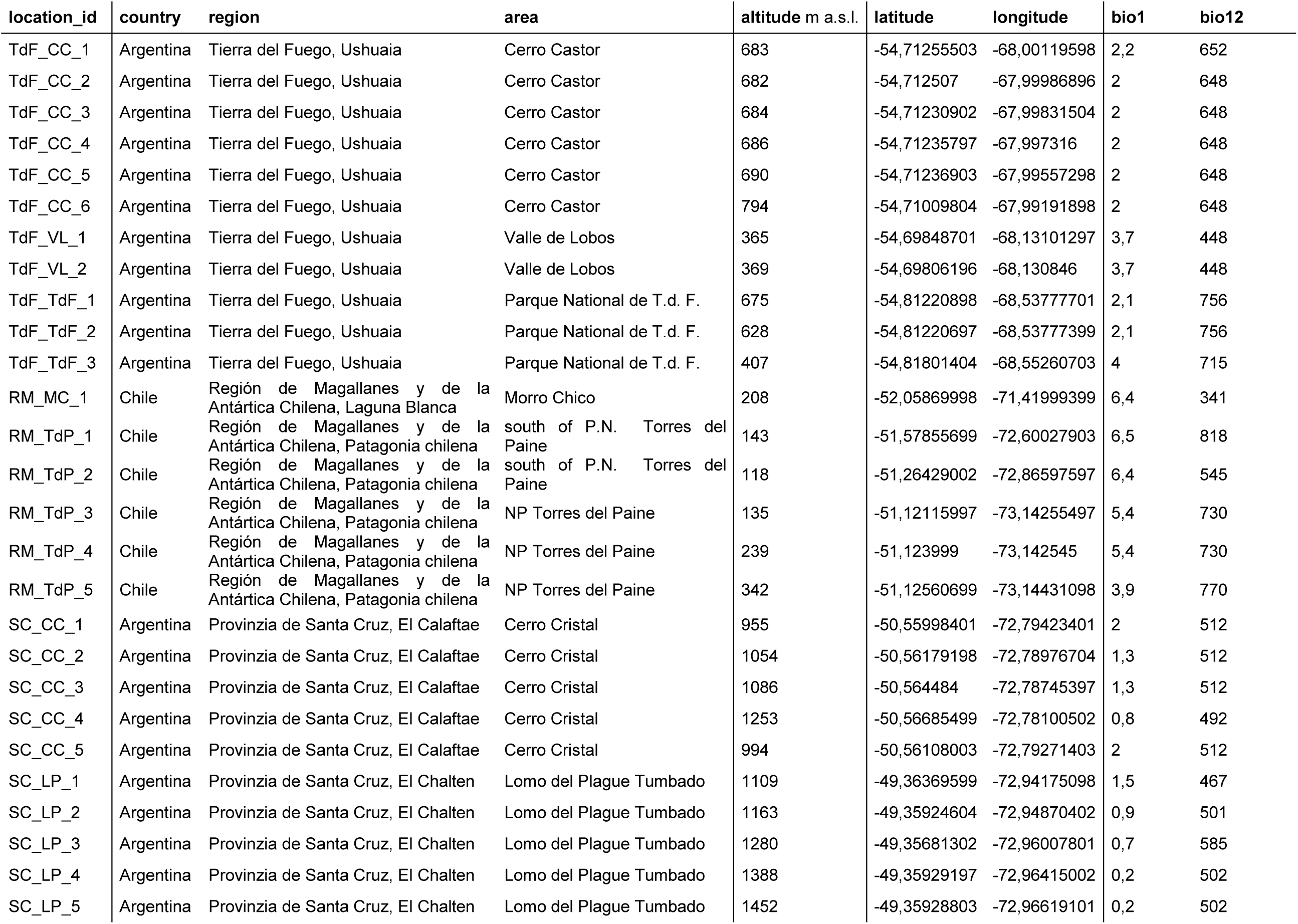
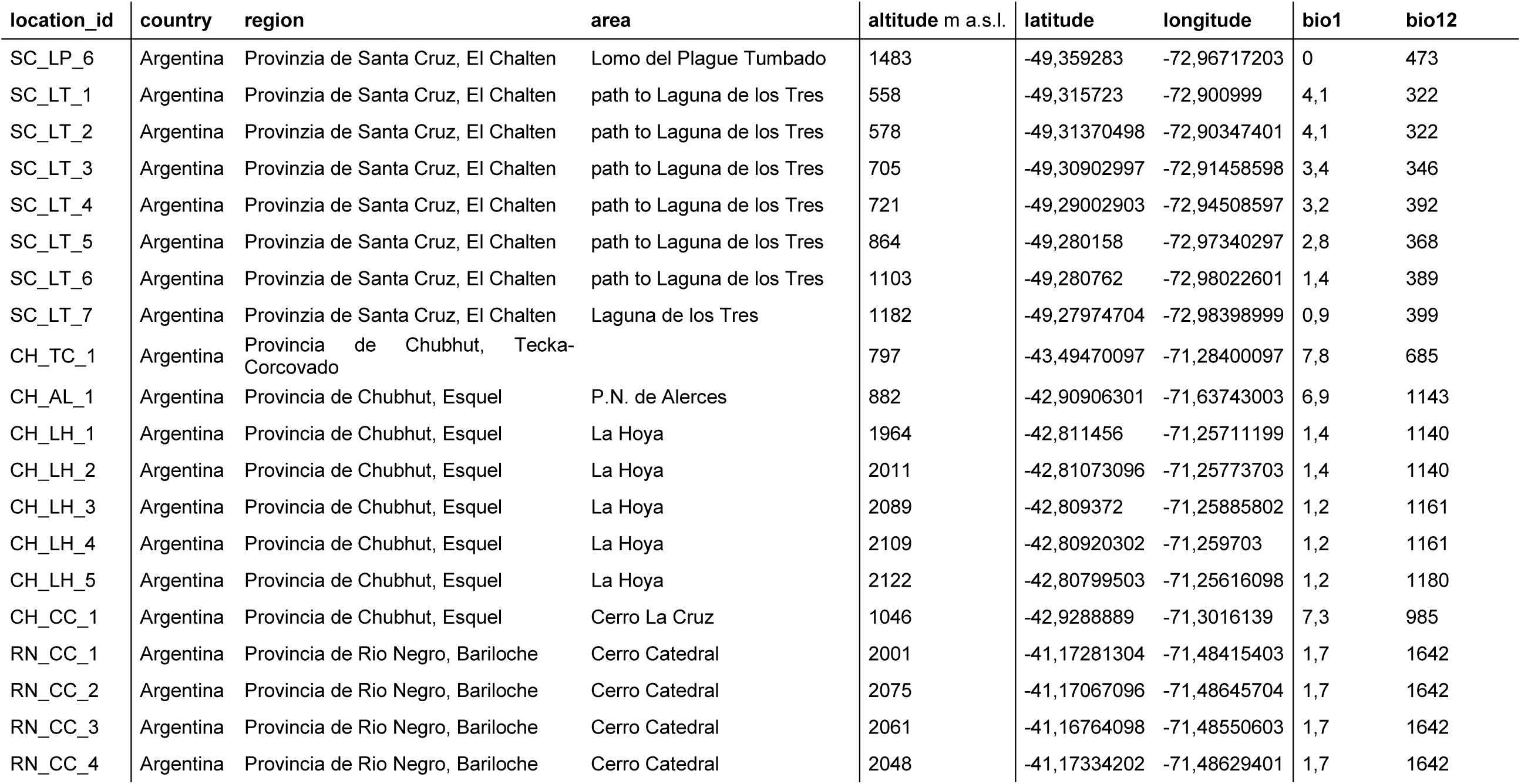
Investigation sites: location_id, location (country, region, area), altitude, latitude, longitude, CHELSA climate variables BIO1 (annual mean temperature) and BIO12 (annual precipitation; Karger *et al*. 2017).

| location_id | country | region | area | altitude m a.s.l. | latitude | longitude | bio1 | bio12 |
| --- | --- | --- | --- | --- | --- | --- | --- | --- |
| TdF_GM_1 | Argentina | Tierra del Fuego, Ushuaia | Glaciar Martial | 570 | -54,78991798 | -68,39034302 | 1,6 | 489 |
| TdF_GM_2 | Argentina | Tierra del Fuego, Ushuaia | Glaciar Martial | 587 | -54,78952697 | -68,39129596 | 1,6 | 489 |
| TdF_GM_3 | Argentina | Tierra del Fuego, Ushuaia | Glaciar Martial | 593 | -54,789 | -68,39313998 | 1,8 | 526 |
| TdF_CB_1 | Chile | Tierra del Fuego, Región de Magallanes y de la Antártica Chilena, Isla Navarino | Cerro Bandera | 607 | -54,96288098 | -67,634564 | 2,7 | 701 |
| TdF_CB_2 | Chile | Tierra del Fuego, Región de Magallanes y de la Antártica Chilena, Isla Navarino | Cerro Bandera | 623 | -54,96410097 | -67,63851698 | 2,7 | 701 |
| TdF_CB_3 | Chile | Tierra del Fuego, Región de Magallanes y de la Antártica Chilena, Isla Navarino | Cerro Bandera | 619 | -54,96530604 | -67,64167897 | 2,7 | 701 |
| TdF_CB_4 | Chile | Tierra del Fuego, Región de Magallanes y de la Antártica Chilena, Isla Navarino | Cerro Bandera | 657 | -54,97040299 | -67,64538897 | 2,3 | 737 |
| TdF_CB_5 | Chile | Tierra del Fuego, Región de Magallanes y de la Antártica Chilena, Isla Navarino | Cerro Bandera | 694 | -54,973166 | -67,642288 | 2,3 | 737 |
| TdF_CB_6 | Chile | Tierra del Fuego, Región de Magallanes y de la Antártica Chilena, Isla Navarino | Cerro Bandera | 702 | -54,97335803 | -67,64681004 | 2,3 | 737 |
| TdF_CB_7 | Chile | Tierra del Fuego, Región de Magallanes y de la Antártica Chilena, Isla Navarino | Cerro Bandera | 688 | -54,97366698 | -67,65027696 | 3,3 | 650 |
| TdF_CB_8 | Chile | Tierra del Fuego, Región de Magallanes y de la Antártica Chilena, Isla Navarino | Cerro Bandera | 680 | -54,97164301 | -67,632565 | 2,6 | 628 |
| TdF_CB_9 | Chile | Tierra del Fuego, Región de Magallanes y de la Antártica Chilena, Isla Navarino | Cerro Bandera | 730 | -54,97210502 | -67,63245997 | 2,6 | 628 |
| TdF_CB_10 | Chile | Tierra del Fuego, Región de Magallanes y de la Antártica Chilena, Isla Navarino | Cerro Bandera | 744 | -54,97210301 | -67,63207901 | 2,6 | 628 |
| TdF_CB_11 | Chile | Tierra del Fuego, Región de Magallanes y de la Antártica Chilena, Isla Navarino | Cerro Bandera | 741 | -54,97349499 | -67,63205403 | 2,6 | 628 |
| TdF_CB_12 | Chile | Tierra del Fuego, Región de Magallanes y de la Antártica Chilena, Isla Navarino | Cerro Bandera | 765 | -54,97333699 | -67,63534997 | 1,9 | 739 |
| TdF_CB_13 | Chile | Tierra del Fuego, Región de Magallanes y de la Antártica Chilena, Isla Navarino | Cerro Bandera | 741 | -54,97261003 | -67,63830702 | 1,9 | 739 |
| TdF_CC_1 | Argentina | Tierra del Fuego, Ushuaia | Cerro Castor | 683 | -54,71255503 | -68,00119598 | 2,2 | 652 |
| TdF_CC_2 | Argentina | Tierra del Fuego, Ushuaia | Cerro Castor | 682 | -54,712507 | -67,99986896 | 2 | 648 |
| TdF_CC_3 | Argentina | Tierra del Fuego, Ushuaia | Cerro Castor | 684 | -54,71230902 | -67,99831504 | 2 | 648 |
| TdF_CC_4 | Argentina | Tierra del Fuego, Ushuaia | Cerro Castor | 686 | -54,71235797 | -67,997316 | 2 | 648 |
| TdF_CC_5 | Argentina | Tierra del Fuego, Ushuaia | Cerro Castor | 690 | -54,71236903 | -67,99557298 | 2 | 648 |
| TdF_CC_6 | Argentina | Tierra del Fuego, Ushuaia | Cerro Castor | 794 | -54,71009804 | -67,99191898 | 2 | 648 |
| TdF_VL_1 | Argentina | Tierra del Fuego, Ushuaia | Valle de Lobos | 365 | -54,69848701 | -68,13101297 | 3,7 | 448 |
| TdF_VL_2 | Argentina | Tierra del Fuego, Ushuaia | Valle de Lobos | 369 | -54,69806196 | -68,130846 | 3,7 | 448 |
| TdF_TdF_1 | Argentina | Tierra del Fuego, Ushuaia | Parque Nacional de T.d. F. | 675 | -54,81220898 | -68,53777701 | 2,1 | 756 |
| TdF_TdF_2 | Argentina | Tierra del Fuego, Ushuaia | Parque Nacional de T.d. F. | 628 | -54,81220697 | -68,53777399 | 2,1 | 756 |
| TdF_TdF_3 | Argentina | Tierra del Fuego, Ushuaia | Parque Nacional de T.d. F. | 407 | -54,81801404 | -68,55260703 | 4 | 715 |
| RM_MC_1 | Chile | Región de Magallanes y de la Antártica Chilena, Laguna Blanca | Morro Chico | 208 | -52,05869998 | -71,41999399 | 6,4 | 341 |
| RM_TdP_1 | Chile | Región de Magallanes y de la Antártica Chilena, Patagonia chilena | south of P.N. Torres del Paine | 143 | -51,57855699 | -72,60027903 | 6,5 | 818 |
| RM_TdP_2 | Chile | Región de Magallanes y de la Antártica Chilena, Patagonia chilena | south of P.N. Torres del Paine | 118 | -51,26429002 | -72,86597597 | 6,4 | 545 |
| RM_TdP_3 | Chile | Región de Magallanes y de la Antártica Chilena, Patagonia chilena | NP Torres del Paine | 135 | -51,12115997 | -73,14255497 | 5,4 | 730 |
| RM_TdP_4 | Chile | Región de Magallanes y de la Antártica Chilena, Patagonia chilena | NP Torres del Paine | 239 | -51,123999 | -73,142545 | 5,4 | 730 |
| RM_TdP_5 | Chile | Región de Magallanes y de la Antártica Chilena, Patagonia chilena | NP Torres del Paine | 342 | -51,12560699 | -73,14431098 | 3,9 | 770 |
| SC_CC_1 | Argentina | Provincia de Santa Cruz, El Calafate | Cerro Cristal | 955 | -50,55998401 | -72,79423401 | 2 | 512 |
| SC_CC_2 | Argentina | Provincia de Santa Cruz, El Calafate | Cerro Cristal | 1054 | -50,56179198 | -72,78976704 | 1,3 | 512 |
| SC_CC_3 | Argentina | Provincia de Santa Cruz, El Calafate | Cerro Cristal | 1086 | -50,564484 | -72,78745397 | 1,3 | 512 |
| SC_CC_4 | Argentina | Provincia de Santa Cruz, El Calafate | Cerro Cristal | 1253 | -50,56685499 | -72,78100502 | 0,8 | 492 |
| SC_CC_5 | Argentina | Provincia de Santa Cruz, El Calafate | Cerro Cristal | 994 | -50,56108003 | -72,79271403 | 2 | 512 |
| SC_LP_1 | Argentina | Provincia de Santa Cruz, El Chalten | Lomo del Plague Tumbado | 1109 | -49,36369599 | -72,94175098 | 1,5 | 467 |
| SC_LP_2 | Argentina | Provincia de Santa Cruz, El Chalten | Lomo del Plague Tumbado | 1163 | -49,35924604 | -72,94870402 | 0,9 | 501 |
| SC_LP_3 | Argentina | Provincia de Santa Cruz, El Chalten | Lomo del Plague Tumbado | 1280 | -49,35681302 | -72,96007801 | 0,7 | 585 |
| SC_LP_4 | Argentina | Provincia de Santa Cruz, El Chalten | Lomo del Plague Tumbado | 1388 | -49,35929197 | -72,96415002 | 0,2 | 502 |
| SC_LP_5 | Argentina | Provincia de Santa Cruz, El Chalten | Lomo del Plague Tumbado | 1452 | -49,35928803 | -72,96619101 | 0,2 | 502 |
| SC_LP_6 | Argentina | Provincia de Santa Cruz, El Chalten | Lomo del Plague Tumbado | 1483 | -49,359283 | -72,96717203 | 0 | 473 |
| SC_LT_1 | Argentina | Provincia de Santa Cruz, El Chalten | path to Laguna de los Tres | 558 | -49,315723 | -72,900999 | 4,1 | 322 |
| SC_LT_2 | Argentina | Provincia de Santa Cruz, El Chalten | path to Laguna de los Tres | 578 | -49,31370498 | -72,90347401 | 4,1 | 322 |
| SC_LT_3 | Argentina | Provincia de Santa Cruz, El Chalten | path to Laguna de los Tres | 705 | -49,30902997 | -72,91458598 | 3,4 | 346 |
| SC_LT_4 | Argentina | Provincia de Santa Cruz, El Chalten | path to Laguna de los Tres | 721 | -49,29002903 | -72,94508597 | 3,2 | 392 |
| SC_LT_5 | Argentina | Provincia de Santa Cruz, El Chalten | path to Laguna de los Tres | 864 | -49,280158 | -72,97340297 | 2,8 | 368 |
| SC_LT_6 | Argentina | Provincia de Santa Cruz, El Chalten | path to Laguna de los Tres | 1103 | -49,280762 | -72,98022601 | 1,4 | 389 |
| SC_LT_7 | Argentina | Provincia de Santa Cruz, El Chalten | Laguna de los Tres | 1182 | -49,27974704 | -72,98398999 | 0,9 | 399 |
| CH_TC_1 | Argentina | Provincia de Chubhut, Tecka-Corcovado |  | 797 | -43,49470097 | -71,28400097 | 7,8 | 685 |
| CH_AL_1 | Argentina | Provincia de Chubhut, Esquel | P.N. de Alerces | 882 | -42,90906301 | -71,63743003 | 6,9 | 1143 |
| CH_LH_1 | Argentina | Provincia de Chubhut, Esquel | La Hoya | 1964 | -42,811456 | -71,25711199 | 1,4 | 1140 |
| CH_LH_2 | Argentina | Provincia de Chubhut, Esquel | La Hoya | 2011 | -42,81073096 | -71,25773703 | 1,4 | 1140 |
| CH_LH_3 | Argentina | Provincia de Chubhut, Esquel | La Hoya | 2089 | -42,809372 | -71,25885802 | 1,2 | 1161 |
| CH_LH_4 | Argentina | Provincia de Chubhut, Esquel | La Hoya | 2109 | -42,80920302 | -71,259703 | 1,2 | 1161 |
| CH_LH_5 | Argentina | Provincia de Chubhut, Esquel | La Hoya | 2122 | -42,80799503 | -71,25616098 | 1,2 | 1180 |
| CH_CC_1 | Argentina | Provincia de Chubhut, Esquel | Cerro La Cruz | 1046 | -42,9288889 | -71,3016139 | 7,3 | 985 |
| RN_CC_1 | Argentina | Provincia de Rio Negro, Bariloche | Cerro Catedral | 2001 | -41,17281304 | -71,48415403 | 1,7 | 1642 |
| RN_CC_2 | Argentina | Provincia de Rio Negro, Bariloche | Cerro Catedral | 2075 | -41,17067096 | -71,48645704 | 1,7 | 1642 |
| RN_CC_3 | Argentina | Provincia de Rio Negro, Bariloche | Cerro Catedral | 2061 | -41,16764098 | -71,48550603 | 1,7 | 1642 |
| RN_CC_4 | Argentina | Provincia de Rio Negro, Bariloche | Cerro Catedral | 2048 | -41,17334202 | -71,48629401 | 1,7 | 1642 |

---

**Table S2a:**
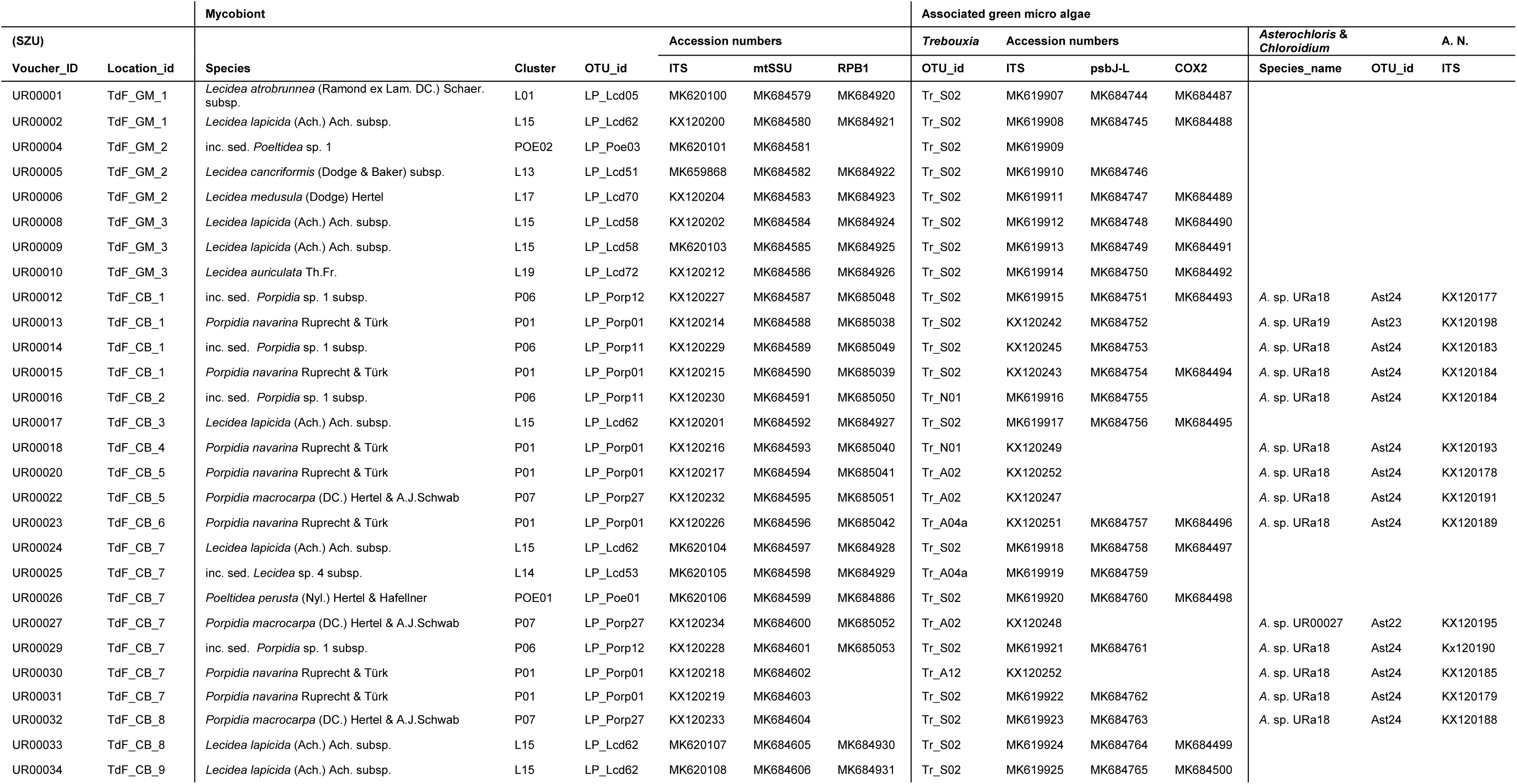
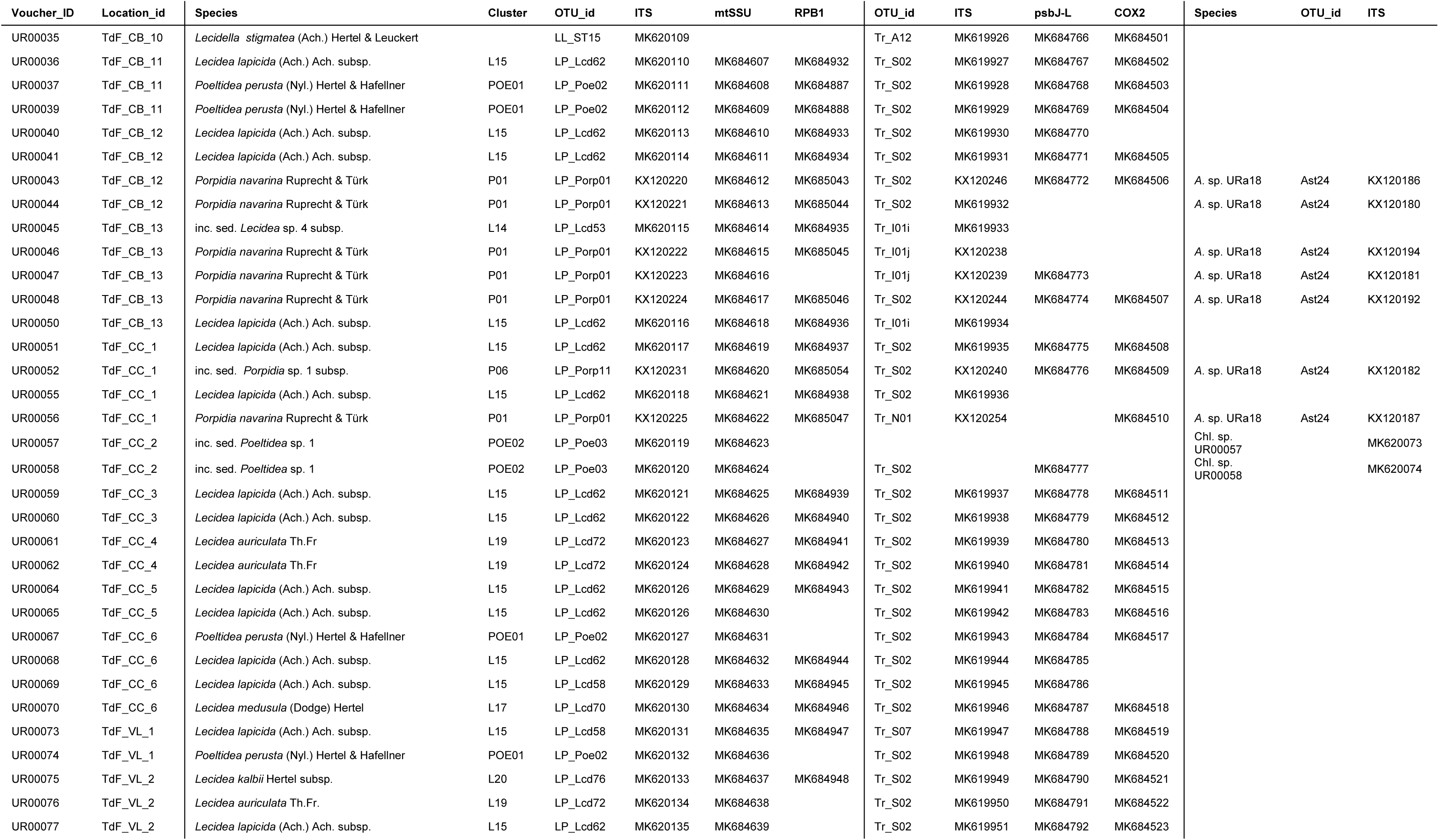
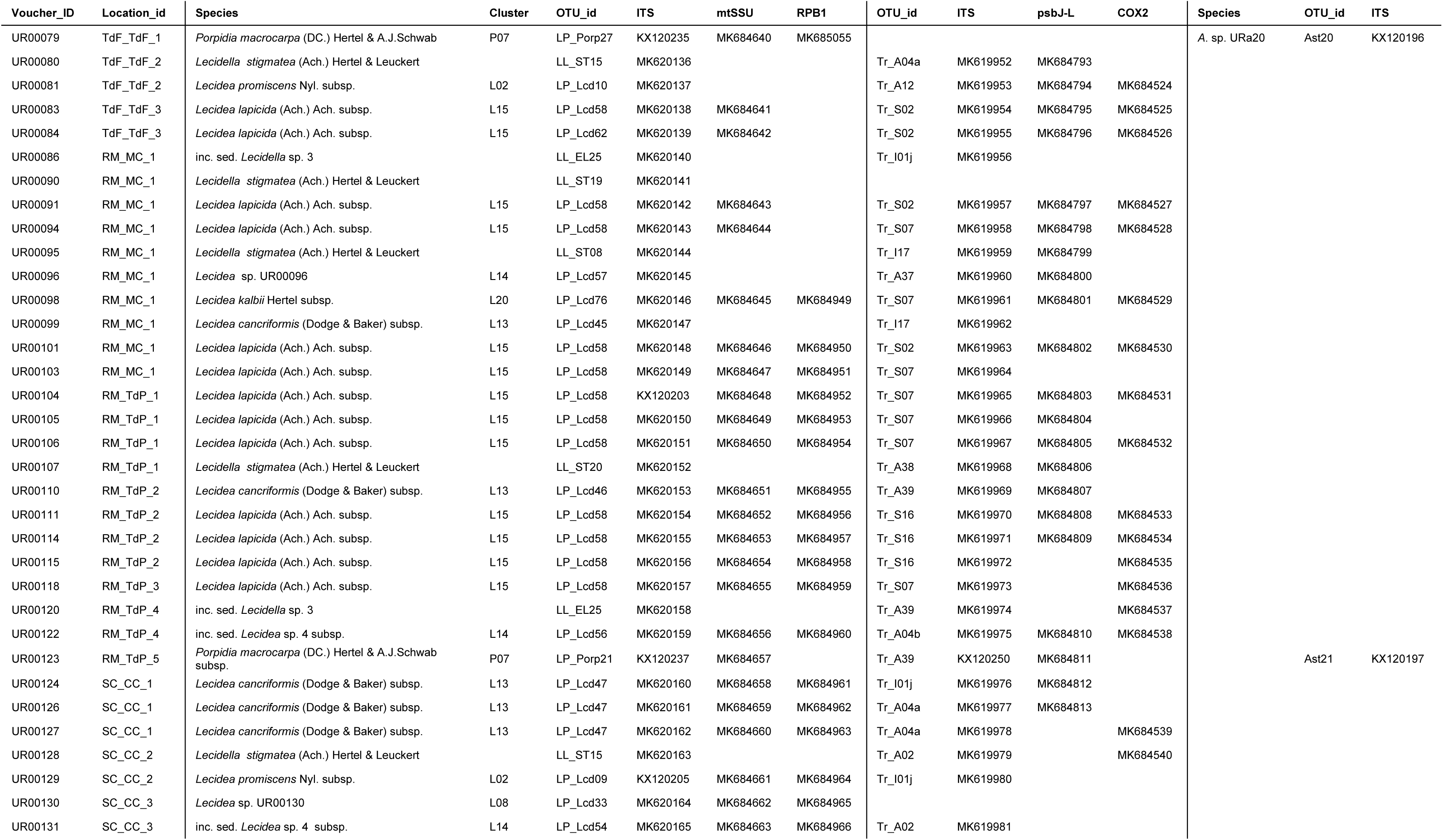
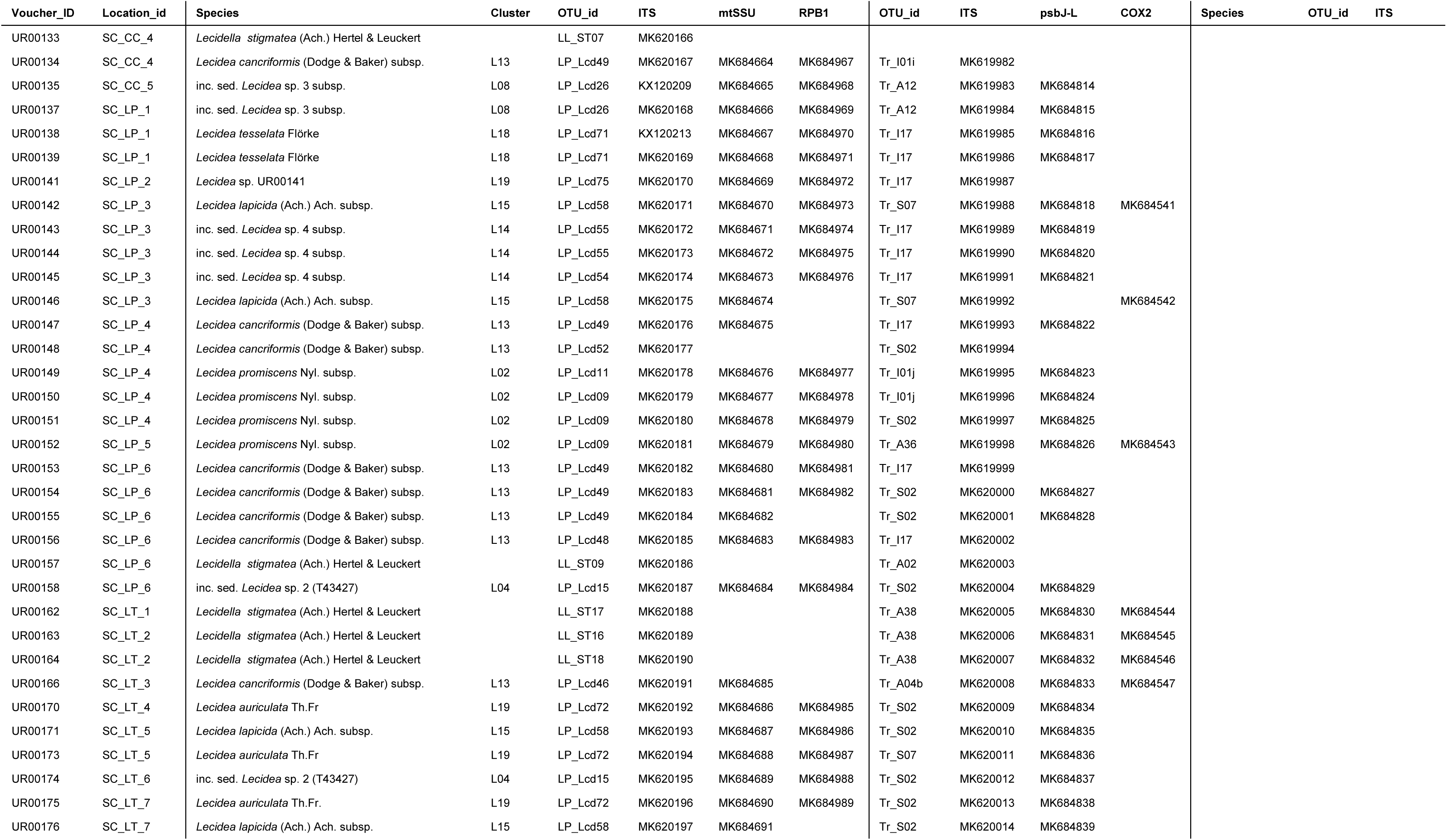
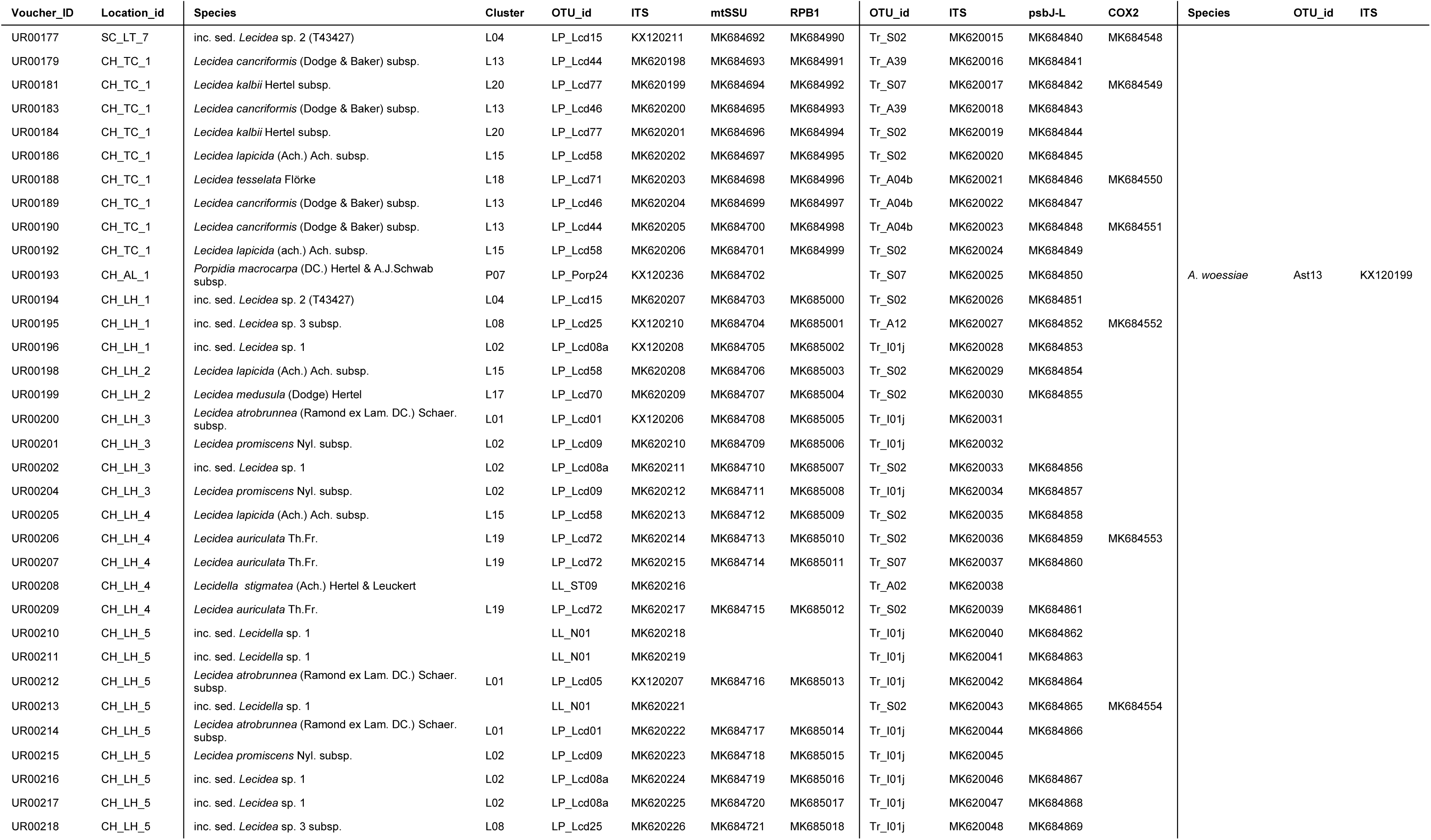
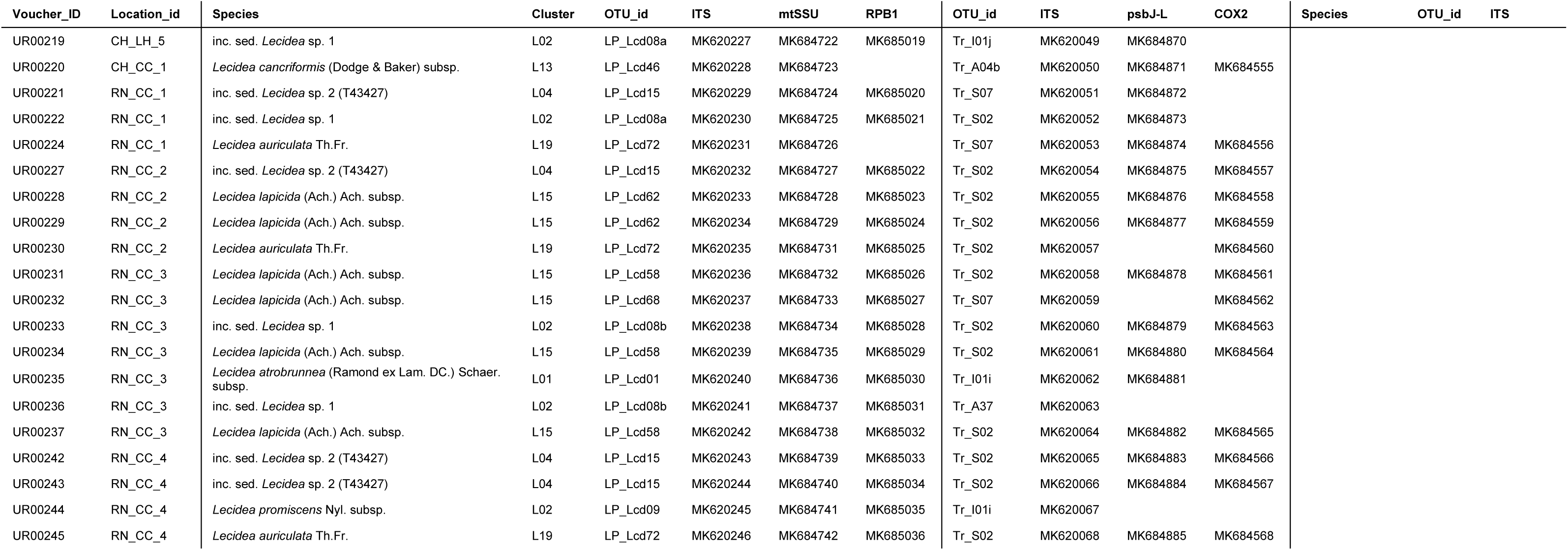
Saxicolous lecideoid lichen specimens from southern South America (sSA) collected by the first author and used in this study. Taxonomic information was taken from the following literature: Ruprecht et al. (2010); Fryday and Hertel (2014); Hafellner and Türk (2016); Ruprecht et al. (2016).

**Table S2b:**
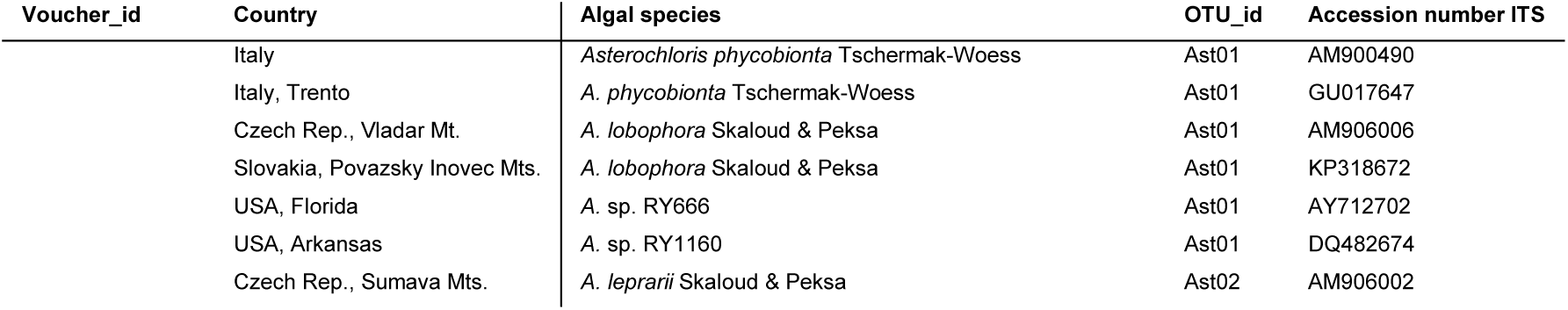
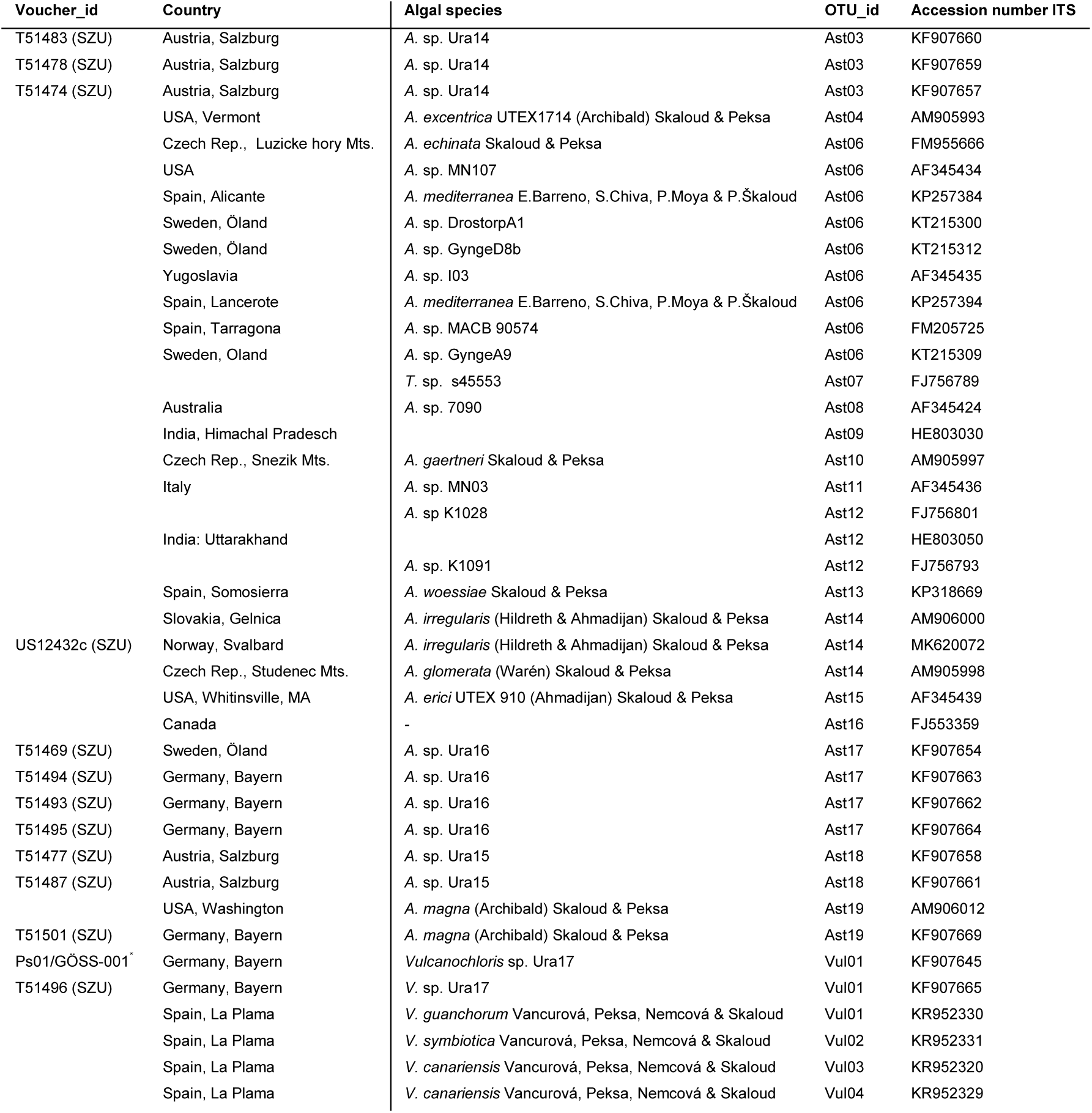
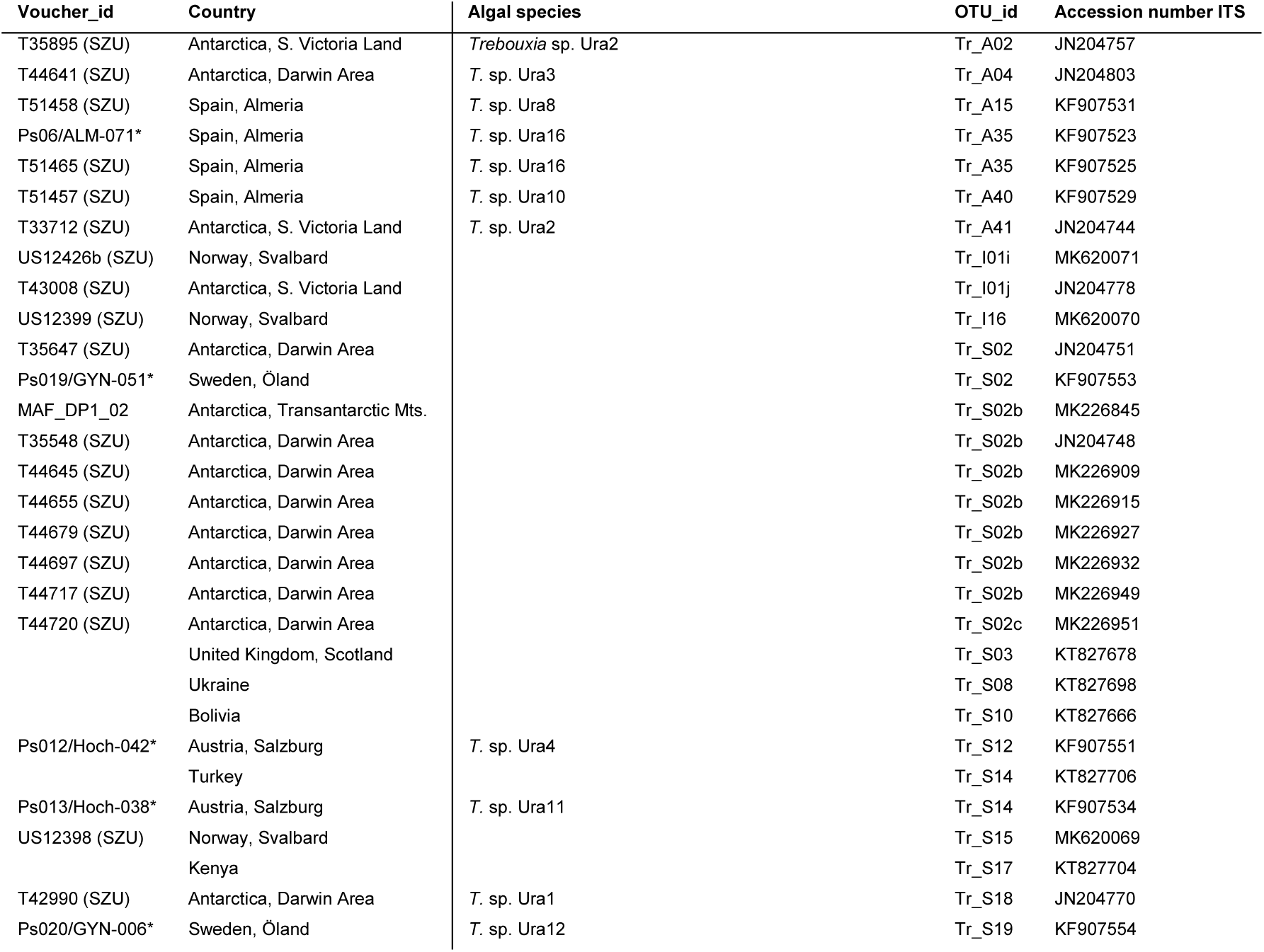
Additional photobiont (*Asterochloris/Vulcanochloris, Trebouxia*) sequences: Newly generated accessions (marked in bold) and downloaded from GenBank (NCBI) worldwide. The representative sequences for the *Trebouxia* OTUs, as defined by Leavitt et al. (2015) that were additionally included to the phylogeny (*Trebouxia* - ITS, Figure 6) are not mentioned in the table. Taxonomic information was taken from the following literature: Leavitt *et al*. (2015); Moya *et al*. (2015); Skaloud *et al*. (2015); Vancurova *et al*. (2015)

| Voucher_id | Country | Algal species | OTU_id | Accession number ITS |
| --- | --- | --- | --- | --- |
|  | Italy | *Asterochloris phycobionta* Tschermak-Woess | Ast01 | AM900490 |
|  | Italy, Trento | *A. phycobionta* Tschermak-Woess | Ast01 | GU017647 |
|  | Czech Rep., Viadar Mt. | *A. lobophora* Skaloud & Peksa | Ast01 | AM906006 |
|  | Slovakia, Povazsky Inovec Mts. | *A. lobophora* Skaloud & Peksa | Ast01 | KP318672 |
|  | USA, Florida | *A. sp.* RY666 | Ast01 | AY712702 |
|  | USA, Arkansas | *A. sp.* RY1160 | Ast01 | DQ482674 |
|  | Czech Rep., Sumava Mts. | *A. leprarii* Skaloud & Peksa | Ast02 | AM906002 |
| T51483 (SZU) | Austria, Salzburg | A. sp. Ura14 | Ast03 | KF907660 |
| T51478 (SZU) | Austria, Salzburg | A. sp. Ura14 | Ast03 | KF907659 |
| T51474 (SZU) | Austria, Salzburg | A. sp. Ura14 | Ast03 | KF907657 |
|  | USA, Vermont | *A. excentrica* UTEX1714 (Archibald) Skaloud & Peksa | Ast04 | AM905993 |
|  | Czech Rep., Luzické hory Mts. | *A. echinata* Skaloud & Peksa | Ast06 | FM955666 |
|  | USA | A. sp. MN107 | Ast06 | AF345434 |
|  | Spain, Alicante | *A. mediterranea* E.Barreno, S.Chiva, P.Moya & P.Škaloud | Ast06 | KP257384 |
|  | Sweden, Öland | A. sp. DrostopA1 | Ast06 | KT215300 |
|  | Sweden, Öland | A. sp. Gynged8b | Ast06 | KT215312 |
|  | Yugoslavia | A. sp. I03 | Ast06 | AF345435 |
|  | Spain, Lancerote | *A. mediterranea* E.Barreno, S.Chiva, P.Moya & P.Škaloud | Ast06 | KP257394 |
|  | Spain, Tarragona | A. sp. MACB 90574 | Ast06 | FM205725 |
|  | Sweden, Öland | A. sp. GyngedA9 | Ast06 | KT215309 |
|  |  | *T. sp.* s45553 | Ast07 | FJ756789 |
|  | Australia | A. sp. 7090 | Ast08 | AF345424 |
|  | India, Himachal Pradesh |  | Ast09 | HE803030 |
|  | Czech Rep., Sněžník Mts. | *A. gaertneri* Skaloud & Peksa | Ast10 | AM905997 |
|  | Italy | A. sp. MN03 | Ast11 | AF345436 |
|  |  | A. sp. K1028 | Ast12 | FJ756801 |
|  | India: Uttarakhand |  | Ast12 | HE803050 |
|  |  | A. sp. K1091 | Ast12 | FJ756793 |
|  | Spain, Somosierra | *A. woessiae* Skaloud & Peksa | Ast13 | KP318669 |
|  | Slovakia, Gelnica | *A. irregularis* (Hildreth & Ahmadjian) Skaloud & Peksa | Ast14 | AM906000 |
| US12432c (SZU) | Norway, Svalbard | *A. irregularis* (Hildreth & Ahmadjian) Skaloud & Peksa | Ast14 | MK620072 |
|  | Czech Rep., Studenec Mts. | *A. glomerata* (Warén) Skaloud & Peksa | Ast14 | AM905998 |
|  | USA, Whitinsville, MA | *A. erici* UTEX 910 (Ahmadjian) Skaloud & Peksa | Ast15 | AF345439 |
|  | Canada | - | Ast16 | FJ553359 |
| T51469 (SZU) | Sweden, Öland | A. sp. Ura16 | Ast17 | KF907654 |
| T51494 (SZU) | Germany, Bayern | A. sp. Ura16 | Ast17 | KF907663 |
| T51493 (SZU) | Germany, Bayern | A. sp. Ura16 | Ast17 | KF907662 |
| T51495 (SZU) | Germany, Bayern | A. sp. Ura16 | Ast17 | KF907664 |
| T51477 (SZU) | Austria, Salzburg | A. sp. Ura15 | Ast18 | KF907658 |
| T51487 (SZU) | Austria, Salzburg | A. sp. Ura15 | Ast18 | KF907661 |
|  | USA, Washington | *A. magna* (Archibald) Skaloud & Peksa | Ast19 | AM906012 |
| T51501 (SZU) | Germany, Bayern | *A. magna* (Archibald) Skaloud & Peksa | Ast19 | KF907669 |
| Ps01/GÖSS-001 | Germany, Bayern | *Vulcanochloris* sp. Ura17 | Vul01 | KF907645 |
| T51496 (SZU) | Germany, Bayern | *V. sp.* Ura17 | Vul01 | KF907665 |
|  | Spain, La Plama | *V. guanchorum* Vancurová, Peksa, Nemcová & Skaloud | Vul01 | KR952330 |
|  | Spain, La Plama | *V. symbiotica* Vancurová, Peksa, Nemcová & Skaloud | Vul02 | KR952331 |
|  | Spain, La Plama | *V. canariensis* Vancurová, Peksa, Nemcová & Skaloud | Vul03 | KR952320 |
|  | Spain, La Plama | *V. canariensis* Vancurová, Peksa, Nemcová & Skaloud | Vul04 | KR952329 |
| T35895 (SZU) | Antarctica, S. Victoria Land | *Trebouxia* sp. Ura2 | Tr_A02 | JN204757 |
| T44641 (SZU) | Antarctica, Darwin Area | *T. sp.* Ura3 | Tr_A04 | JN204803 |
| T51458 (SZU) | Spain, Almeria | *T. sp.* Ura8 | Tr_A15 | KF907531 |
| Ps06/ALM-071* | Spain, Almeria | *T. sp.* Ura16 | Tr_A35 | KF907523 |
| T51465 (SZU) | Spain, Almeria | *T. sp.* Ura16 | Tr_A35 | KF907525 |
| T51457 (SZU) | Spain, Almeria | *T. sp.* Ura10 | Tr_A40 | KF907529 |
| T33712 (SZU) | Antarctica, S. Victoria Land | *T. sp.* Ura2 | Tr_A41 | JN204744 |
| US12426b (SZU) | Norway, Svalbard |  | Tr_I01i | MK620071 |
| T43008 (SZU) | Antarctica, S. Victoria Land |  | Tr_I01j | JN204778 |
| US12399 (SZU) | Norway, Svalbard |  | Tr_I16 | MK620070 |
| T35647 (SZU) | Antarctica, Darwin Area |  | Tr_S02 | JN204751 |
| Ps019/GYN-051* | Sweden, Öland |  | Tr_S02 | KF907553 |
| MAF_DP1_02 | Antarctica, Transantarctic Mts. |  | Tr_S02b | MK226845 |
| T35548 (SZU) | Antarctica, Darwin Area |  | Tr_S02b | JN204748 |
| T44645 (SZU) | Antarctica, Darwin Area |  | Tr_S02b | MK226909 |
| T44655 (SZU) | Antarctica, Darwin Area |  | Tr_S02b | MK226915 |
| T44679 (SZU) | Antarctica, Darwin Area |  | Tr_S02b | MK226927 |
| T44697 (SZU) | Antarctica, Darwin Area |  | Tr_S02b | MK226932 |
| T44717 (SZU) | Antarctica, Darwin Area |  | Tr_S02b | MK226949 |
| T44720 (SZU) | Antarctica, Darwin Area |  | Tr_S02c | MK226951 |
|  | United Kingdom, Scotland |  | Tr_S03 | KT827678 |
|  | Ukraine |  | Tr_S08 | KT827698 |
|  | Bolivia |  | Tr_S10 | KT827666 |
| Ps012/Hoch-042* | Austria, Salzburg | *T. sp.* Ura4 | Tr_S12 | KF907551 |
|  | Turkey |  | Tr_S14 | KT827706 |
| Ps013/Hoch-038* | Austria, Salzburg | *T. sp.* Ura11 | Tr_S14 | KF907534 |
| US12398 (SZU) | Norway, Svalbard |  | Tr_S15 | MK620069 |
|  | Kenya |  | Tr_S17 | KT827704 |
| T42990 (SZU) | Antarctica, Darwin Area | *T. sp.* Ura1 | Tr_S18 | JN204770 |
| Ps020/GYN-006* | Sweden, Öland | *T. sp.* Ura12 | Tr_S19 | KF907554 |

**Table 2c:**
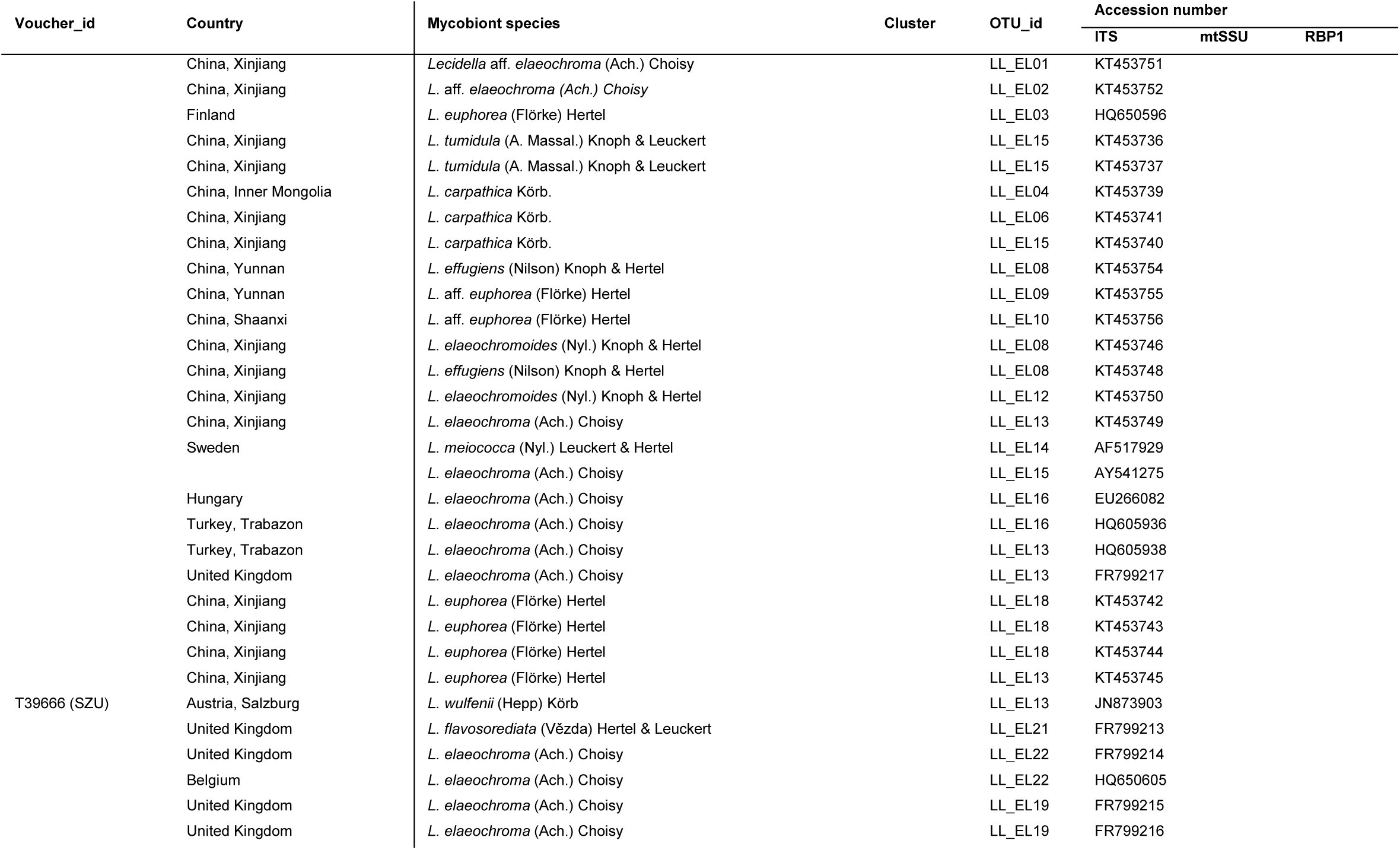
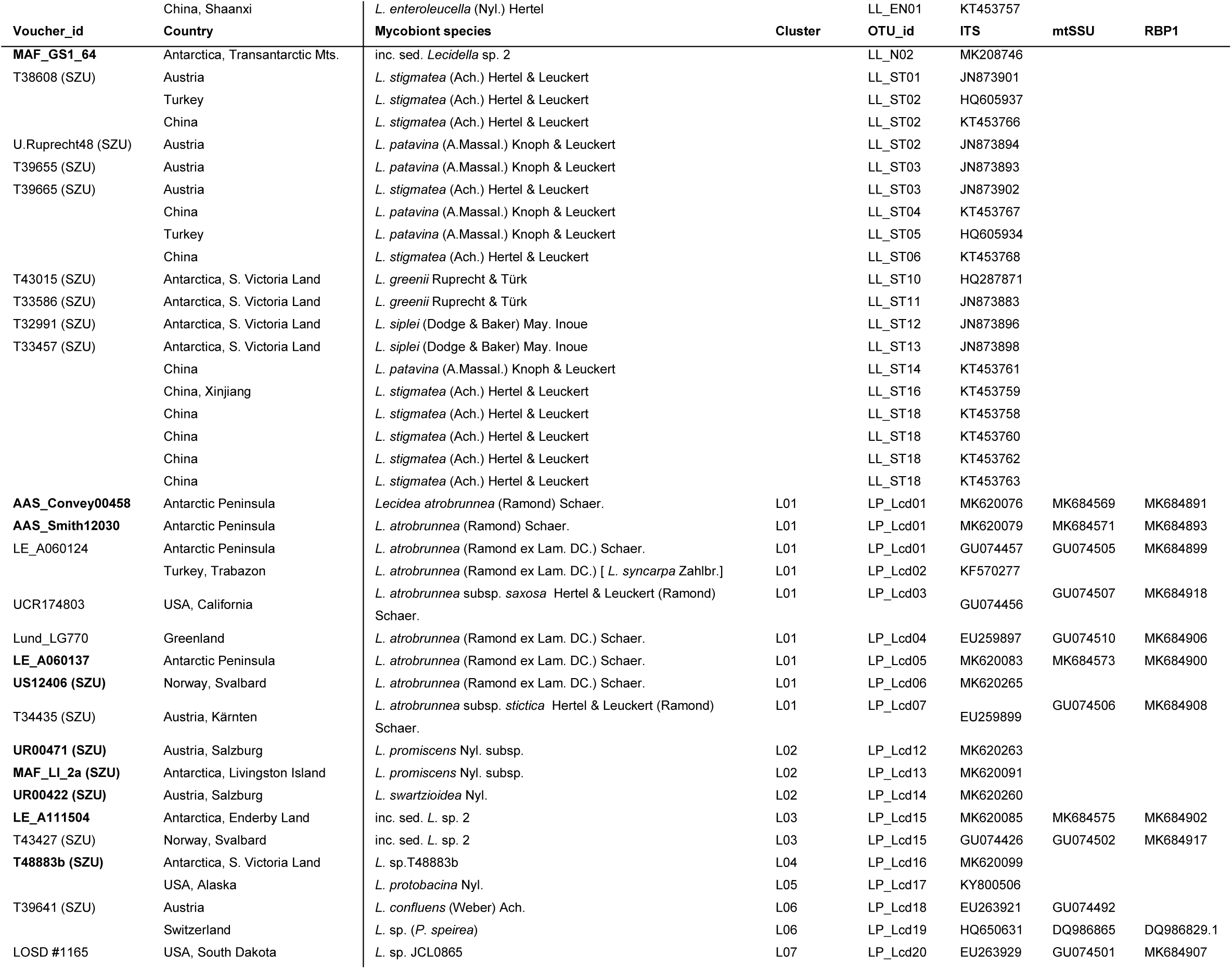
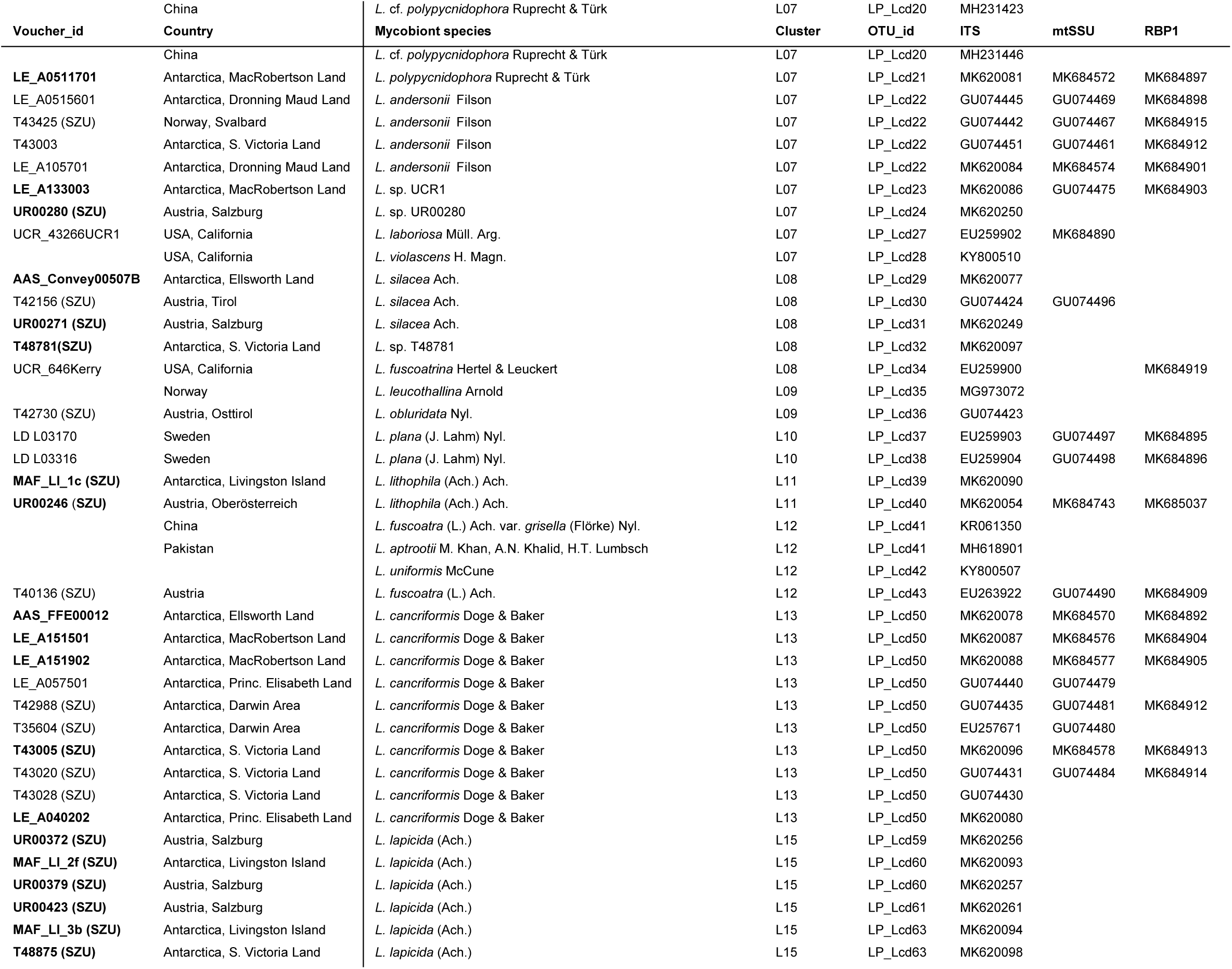
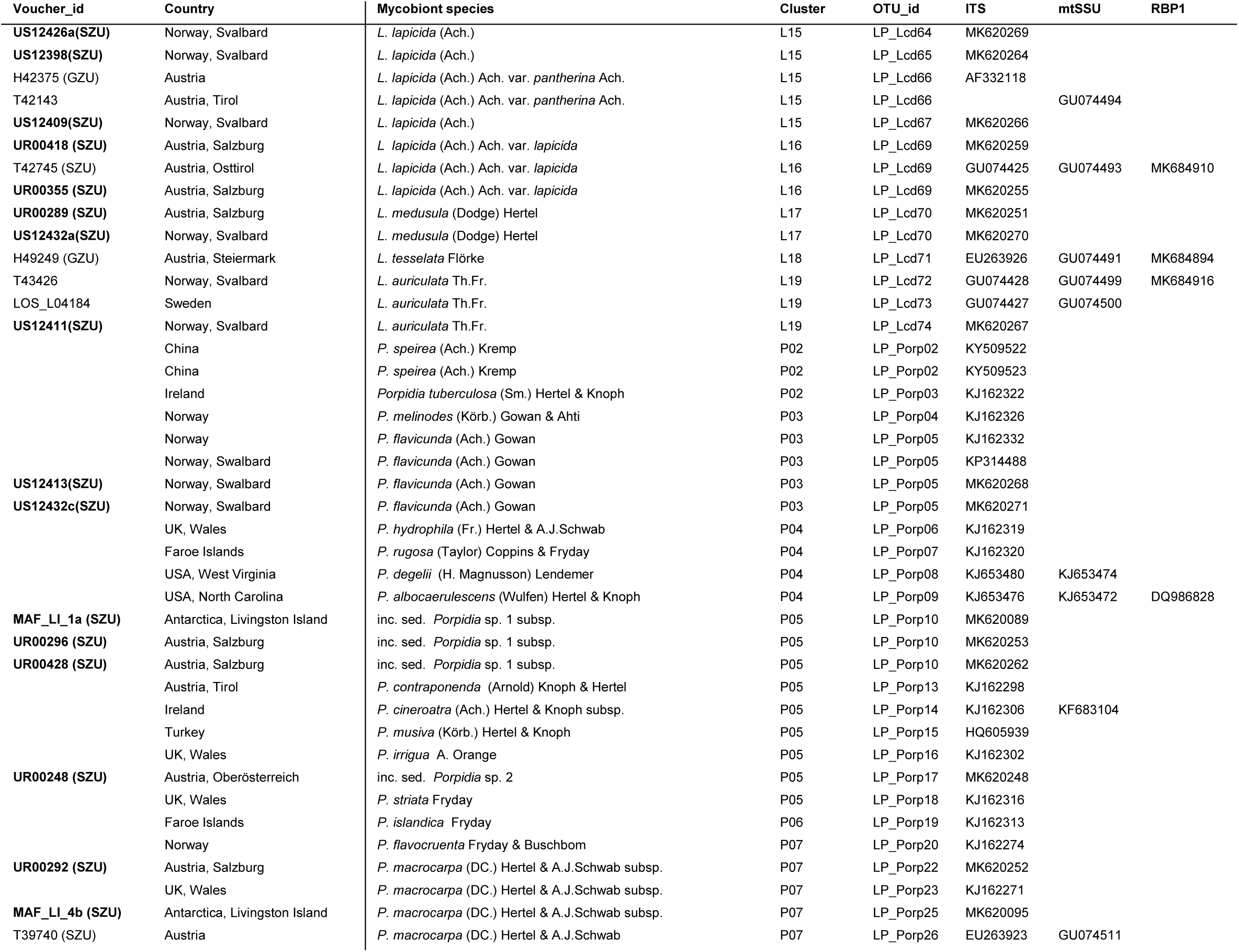
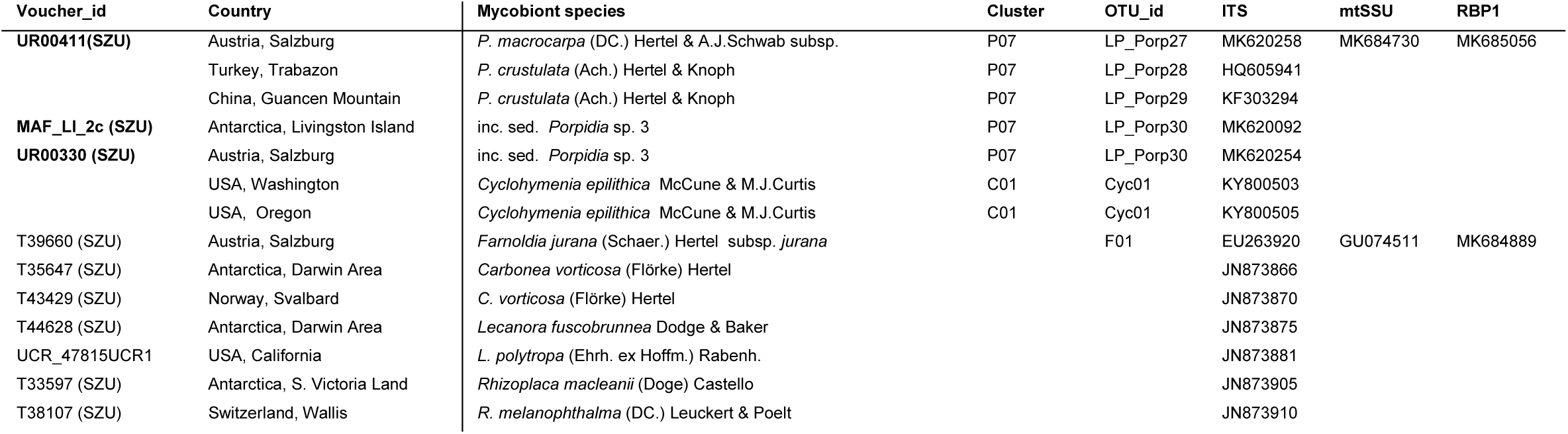
Additional Mycobiont (*Lecidella, Lecidea, Porpidia*) sequences: Own and newly generated (marked in bold) accessions and from Genbank (NCBI) worldwide. Taxonomic information was taken from the following literature: Ruprecht et al. (2010, 2012); Wirth et al. (2013); Orange (2014); Zhao et al. (2015); Hafellner and Türk (2016); McCune et al. (2017).

**Table 2d:**
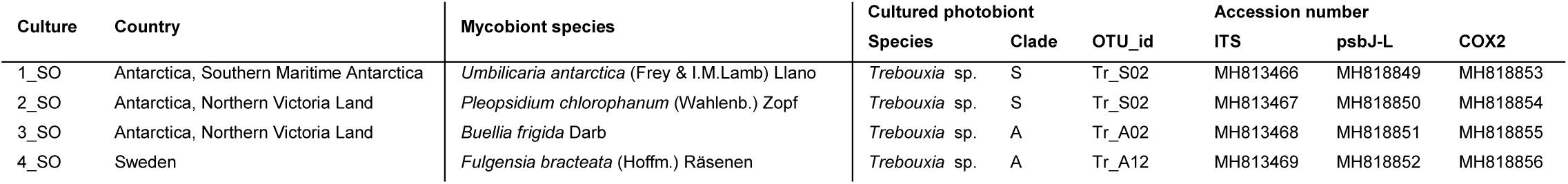
Cultured Trebouxia specimen provided by Sieglinde Ott, Heinrich Heine Universität, Düsseldorf, Germany

**Table S3:**
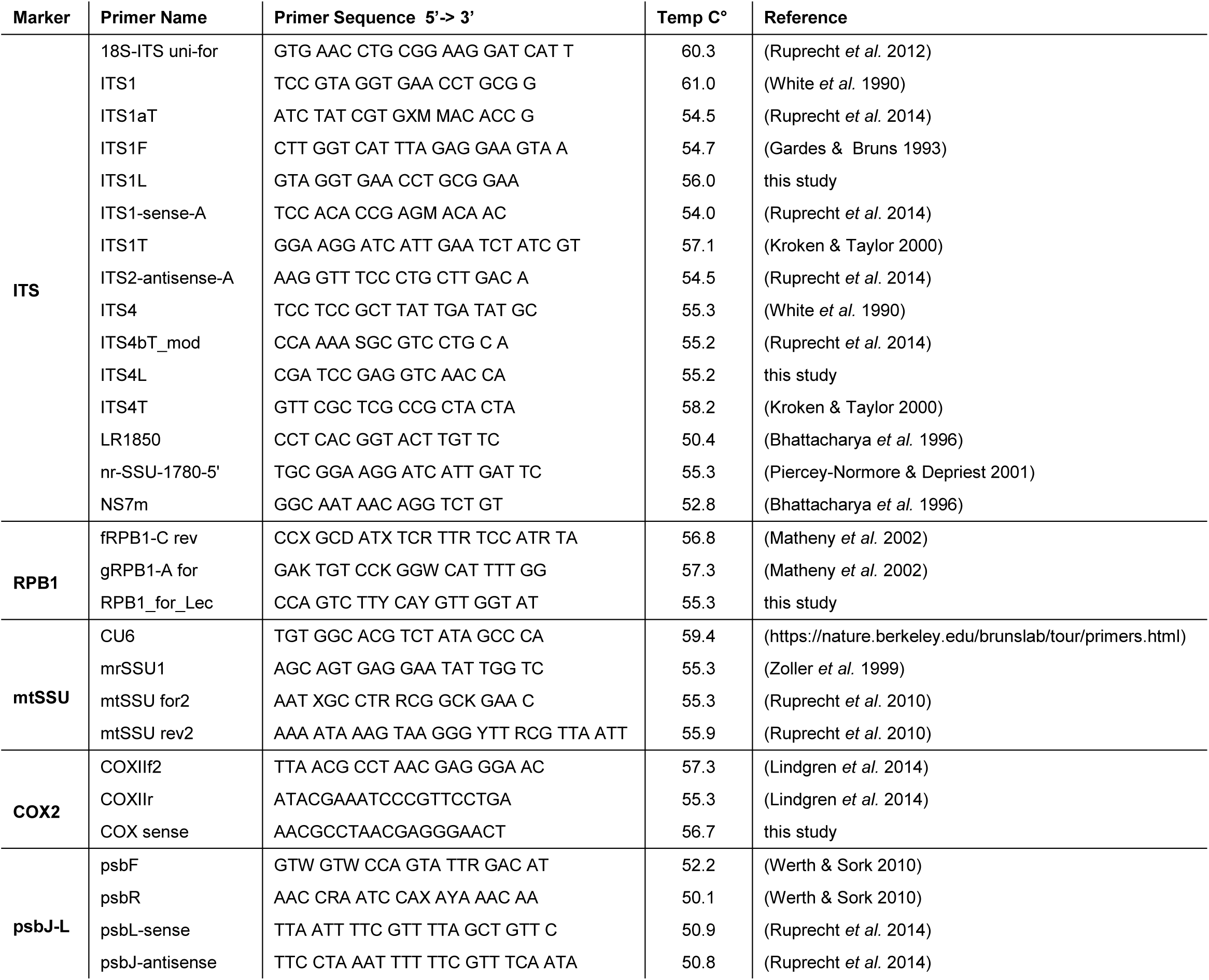
Marker and primer used in this study.

**Table S4:** Evolutionary models of the Maximum Likelihood analyses.

| Phylogeny | Mycobiont | Photobiont |
| --- | --- | --- |
| *Lecidea/Porpidia/Poeltidea* - ITS | TIM2+F+I+G4 |  |
| *Lecidea/Porpidia/Poeltidea* - ITS/mtSSU/RPB1 | TIM2e+I+G4 |  |
| *Lecidella* - ITS | TNe+I+G4 |  |
| *Asterochloris* - ITS |  | TIM2e+I+G4 |
| *Trebouxia*- ITS |  | SYM+I+G4 |
| *Trebouxia* - ITS/psbJ-L/COX2 |  | TPM3u+F+G4 |

**Table S5:** Scores of sequence similarity in % of more than 3 accessions per OTU

| OTU $\geq 3$ accessions | sub - OTU | sequence similarity % |
| --- | --- | --- |
| LP_Lcd01 |  | 99.8 |
| LP_Lcd08 total |  | 97.3 |
|  | LP_08a | 97.9 |
|  | LP_08b | 99.8 |
| LP_Lcd09 |  | 99.4 |
| LP_Lcd15 |  | 98.6 |
| LP_Lcd46 |  | 99.1 |
| LP_Lcd49 |  | 99.2 |
| LP_Lcd50 |  | 98.6 |
| LP_Lcd58 |  | 98.7 |
| LP_Lcd62 |  | 98.9 |
| LP_Lcd71 |  | 98.8 |
| LP_Lcd72 |  | 98.9 |
| LP_Porp 27 |  | 98.5 |
| LL_N01 |  | 99.4 |
| Ast24 |  | 99.8 |
| Tr_A02 |  | 99.4 |
| Tr_A04 total |  | 96.7 |
|  | Tr_A04a | 97.7 |
|  | Tr_A04b | 98.8 |
| Tr_S02 total |  | 94.8 |
|  | Tr_S02 | 98.0 |
|  | Tr_S02b | 98.4 |
|  | Tr_I01i | 97.5 |
|  | Tr_I01j | 98.6 |
| Tr_I17 |  | 99.1 |

